## Supplementary Figure 1 for "Identification of novel PfEMP1 variants containing domain cassettes 11, 15 and 8 that mediate the *Plasmodium falciparum* virulence-associated rosetting phenotype"

| | | | | | | | | | |
|---|---|---|---|---|---|---|---|---|---|
| PFKE01.g2 | DBLα1.1 | CIDRα1.7 | DBLβ1 | DBLγ6 | DBLδ5 | CIDRβ3 | DBLβ6 | DBLγ2 | |
| PFKE01.g6 | DBLα1.1 | CIDRα1.7 | DBLβ1 | DBLβ6 | DBLδ1 | CIDRβ1 | | | |
| PFKE01.g232 | DBLα1.1 | CIDRα1.2 | DBLβ11 | DBLγ1 | DBLε1 | DBLγ8 | DBLζ1 | DBLε5 | |
| PFKE01.g198 | DBLα1.2 | CIDRα1.4 | DBLβ3 | DBLγ11 | DBLγ6 | DBLδ1 | CIDRβ6 | | |
| PFKE01.g1 | DBLα1.4 | CIDRα1.1 | DBLζ3 | DBLγ12 | DBLδ5 | CIDRβ3 | DBLβ6 | DBLγ12 | |
| PFKE01.g197 | DBLα1.5 | CIDRβ4 | DBLβ7 | DBLγ12 | DBLδ1 | CIDRβ3 | | | |
| PFKE01.g5 | DBLα1.7 | CIDRα1.4 | DBLγ11 | DBLβ6 | DBLγ11 | DBLγ2 | | | |
| PFKE01.g9 | DBLα0.1 | CIDRα3.2 | DBLδ1 | CIDRβ5 | | | | | |
| PFKE01.g190 | DBLα0.1 | CIDRα3.1 | DBLδ1 | CIDRβ5 | | | | | |
| PFKE01.g229 | DBLα0.1 | CIDRα3.2 | DBLδ1 | CIDRβ1 | | | | | |
| PFKE01.g282 | DBLα0.1 | CIDRα3.1 | DBLδ1 | CIDRβ1 | | | | | |
| PFKE01.g284 | DBLα0.1 | CIDRα3.2 | DBLδ1 | CIDRβ1 | | | | | |
| PFKE01.g199 | DBLα0.1 | CIDRα2.2 | DBLδ1 | CIDRγ7 | | | | | |
| PFKE01.g27 | DBLα0.11 | CIDRα2.4 | DBLδ1 | CIDRβ1 | | | | | |
| PFKE01.g326 | DBLα0.11 | CIDRα2.4 | DBLδ1 | CIDRβ1 | | | | | |
| PFKE01.g206 | DBLα0.12 | CIDRα2.1 | DBLγ9 | | | | | | |
| PFKE01.g228 | DBLα0.12 | CIDRα2.2 | DBLδ1 | CIDRγ1 | | | | | |
| PFKE01.g283 | DBLα0.13 | CIDRα2.6 | DBLδ1 | CIDRβ6 | | | | | |
| PFKE01.g18 | DBLα0.14 | CIDRα6 | DBLδ1 | CIDRβ5 | | | | | |
| PFKE01.g302 | DBLα0.14 | CIDRα4 | DBLδ1 | CIDRβ1 | | | | | |
| PFKE01.g17 | DBLα0.16 | CIDRα3.4 | DBLδ1 | CIDRβ1 | | | | | |
| PFKE01.g451 | DBLα0.16 | CIDRα3.4 | DBLδ1 | CIDRβ1 | | | | | |
| PFKE01.g19 | DBLα0.17 | CIDRα3.3 | DBLδ1 | CIDRβ1 | | | | | |
| PFKE01.g14 | DBLα0.18 | CIDRα4 | DBLβ5 | DBLγ3 | DBLζ4 | | | | |
| PFKE01.g155 | DBLα0.18 | CIDRα6 | DBLβ4 | DBLγ10 | DBLδ5 | CIDRβ4 | | | |
| PFKE01.g191 | DBLα0.18 | CIDRα6 | DBLβ5 | DBLδ8 | CIDRβ2 | | | | |
| PFKE01.g281 | DBLα0.18 | CIDRα5 | DBLβ5 | DBLγ10 | DBLδ1 | CIDRβ1 | | | |
| PFKE01.g231 | DBLα0.19 | CIDRα2.9 | DBLδ1 | CIDRβ1 | | | | | |
| PFKE01.g388 | DBLα0.2 | CIDRα3.3 | DBLδ1 | CIDRβ1 | | | | | |
| PFKE01.g227 | DBLα0.21 | CIDRα6 | DBLβ5 | DBLγ12 | DBLδ4 | CIDRβ4 | DBLε2 | DBLε7 | DBLε3 |
| PFKE01.g230 | DBLα0.22 | CIDRα3.2 | DBLβ5 | DBLδ1 | CIDRβ1 | | | | |
| PFKE01.g200 | DBLα0.23 | CIDRα5 | DBLβ5 | DBLδ1 | CIDRγ5 | DBLγ3 | DBLζ4 | | |
| PFKE01.g16 | DBLα0.24 | CIDRα3.3 | DBLδ1 | CIDRβ4 | | | | | |
| PFKE01.g195 | DBLα0.3 | CIDRα3.4 | DBLβ5 | DBLγ10 | DBLδ4 | CIDRγ6 | | | |
| PFKE01.g13 | DBLα0.4 | CIDRα6 | DBLδ1 | CIDRγ5 | | | | | |
| PFKE01.g23 | DBLα0.4 | CIDRα3.1 | DBLδ1 | CIDRβ6 | | | | | |
| PFKE01.g7 | DBLα0.5 | CIDRα2.8 | DBLβ8 | DBLδ1 | CIDRβ6 | | | | |
| PFKE01.g21 | DBLα0.5 | CIDRα2.6 | DBLδ1 | CIDRβ6 | | | | | |
| PFKE01.g467 | DBLα0.5 | CIDRα2.9 | DBLδ1 | CIDRβ1 | | | | | |
| PFKE01.g22 | DBLα0.6 | CIDRα3.1 | DBLδ1 | CIDRβ5 | | | | | |
| PFKE01.g45 | DBLα0.6 | CIDRα3.1 | DBLδ8 | CIDRβ4 | | | | | |
| PFKE01.g15 | DBLα0.7 | CIDRα2.2 | DBLδ1 | CIDRβ1 | | | | | |
| PFKE01.g292 | DBLα0.8 | CIDRα2.2 | DBLδ1 | CIDRβ5 | | | | | |
| PFKE01.g10 | DBLα0.9 | CIDRα2.1 | DBLδ1 | CIDRβ1 | | | | | |
| PFKE01.g430 | DBLα0.9 | CIDRα2.1 | DBLδ1 | CIDRγ4 | | | | | |
| PFKE01.g12 | DBLpam1 | DBLpam2 | CIDRpam | DBLpam3 | DBLεpam4 | DBLεpam5 | DBLε10 | | |
| PFKE01.g345 | CIDRβ4 | | | | | | | | |
| PFKE01.g4 | DBLβ13 | DBLβ5 | DBLβ8 | DBLδ1 | CIDRβ1 | | | | |
| PFKE01.g46 | DBLδ1 | CIDRβ5 | | | | | | | |

**Rosetting-associated head structure**

**DBLα1.5 /6 /8** **CIDRβ /γ /δ**

### PFKE01 (9106)

| | | | | | | | | |
|---|---|---|---|---|---|---|---|---|
| PFKE02.g224 | DBLα1.1 | CIDRα1.2 | DBLβ11 | DBLγ1 | DBLε1 | DBLγ8 | DBLζ2 | DBLε5 |
| PFKE02.g1 | DBLα1.2 | CIDRα1.4 | DBLβ1 | DBLβ6 | DBLγ6 | DBLγ11 | DBLδ1 | CIDRβ1 |
| PFKE02.g7 | DBLα1.2 | CIDRα1.7 | DBLγ2 | DBLγ2 | DBLγ4 | DBLζ3 | DBLε6 | |
| PFKE02.g225 | DBLα1.2 | CIDRα1.7 | DBLβ3 | DBLγ11 | DBLδ3 | CIDRγ2 | DBLζ6 | DBLε6 |
| PFKE02.g252 | DBLα1.2 | CIDRα1.5 | DBLγ12 | DBLδ5 | CIDRβ4 | DBLβ6 | DBLγ11 | DBLε4 |
| PFKE02.g253 | DBLα1.2 | CIDRα1.5 | DBLβ3 | DBLγ11 | DBLδ1 | CIDRβ3 | | |
| PFKE02.g24 | DBLα1.3 | DBLε6 | | | | | | |
| PFKE02.g12 | DBLα1.4 | CIDRα1.6 | DBLβ3 | DBLγ10 | DBLδ6 | CIDRβ3 | | |
| PFKE02.g264 | DBLα1.5 | CIDRβ4 | DBLβ7 | DBLγ10 | DBLδ3 | CIDRγ2 | DBLζ6 | DBLε6 |
| PFKE02.g562 | DBLα1.5 | CIDRδ1 | DBLγ11 | DBLγ10 | DBLδ1 | CIDRβ5 | | |
| PFKE02.g6 | DBLα1.6 | CIDRγ3 | DBLγ7 | DBLδ5 | CIDRβ4 | DBLβ9 | DBLγ11 | |
| PFKE02.g3 | DBLα1.7 | CIDRα1.5 | DBLγ12 | DBLδ4 | CIDRγ1 | DBLβ9 | | |
| PFKE02.g4 | DBLα1.7 | CIDRα1.7 | DBLβ3 | DBLγ11 | DBLγ9 | DBLδ1 | CIDRβ1 | |
| PFKE02.g50 | DBLα1.7 | CIDRγ3 | DBLβ6 | DBLγ10 | DBLδ4 | CIDRγ2 | | |
| PFKE02.g88 | DBLα1.7 | CIDRα1.7 | DBLβ12 | DBLγ4 | DBLγ2 | DBLδ1 | CIDRβ5 | |
| PFKE02.g428 | DBLα2 | CIDRα1.1 | DBLβ12 | DBLγ4 | DBLδ1 | CIDRβ5 | | |
| PFKE02.g11 | DBLα0.1 | CIDRα3.2 | DBLδ1 | CIDRβ1 | | | | |
| PFKE02.g254 | DBLα0.1 | CIDRα3.2 | DBLδ1 | CIDRβ1 | | | | |
| PFKE02.g255 | DBLα0.1 | CIDRα3.2 | DBLδ1 | CIDRβ5 | | | | |
| PFKE02.g355 | DBLα0.1 | CIDRα3.1 | DBLδ1 | CIDRγ11 | | | | |
| PFKE02.g508 | DBLα0.1 | CIDRα3.1 | DBLδ1 | CIDRβ1 | | | | |
| PFKE02.g561 | DBLα0.1 | CIDRα3.2 | DBLδ1 | CIDRβ5 | | | | |

**Rosetting-associated head structure**

**DBLα1.5 /6 /8** **CIDRβ /γ /δ**

#### PFKE02 (9626)

| | | | | | | | | |
|---|---|---|---|---|---|---|---|---|
| PFKE02.g38 | DBLα0.1 | CIDRα2.2 | DBLδ1 | CIDRβ1 | | | | |
| PFKE02.g259 | DBLα0.1 | CIDRα2.2 | DBLδ1 | CIDRβ1 | | | | |
| PFKE02.g16 | DBLα0.11 | CIDRα2.4 | DBLδ1 | CIDRβ1 | | | | |
| PFKE02.g258 | DBLα0.11 | CIDRα2.4 | DBLβ5 | DBLδ1 | CIDRβ7 | | | |
| PFKE02.g265 | DBLα0.11 | CIDRα2.1 | DBLδ6 | | | | | |
| PFKE02.g248 | DBLα0.12 | CIDRα2.2 | DBLδ1 | CIDRγ12 | DBLγ13 | DBLζ1 | DBLε6 | |
| PFKE02.g176 | DBLα0.13 | CIDRα2.9 | DBLδ1 | CIDRγ1 | | | | |
| PFKE02.g268 | DBLα0.13 | CIDRα2.5 | DBLδ1 | CIDRγ9 | | | | |
| PFKE02.g17 | DBLα0.14 | CIDRα4 | DBLδ1 | CIDRγ4 | | | | |
| PFKE02.g251 | DBLα0.15 | CIDRα3.2 | DBLδ1 | CIDRγ11 | | | | |
| PFKE02.g262 | DBLα0.15 | CIDRα3.2 | | | | | | |
| PFKE02.g332 | DBLα0.15 | CIDRα3.2 | DBLβ5 | DBLγ11 | DBLδ1 | CIDRβ1 | | |
| PFKE02.g308 | DBLα0.16 | CIDRα3.4 | DBLδ1 | CIDRβ1 | | | | |
| PFKE02.g354 | DBLα0.18 | CIDRα5 | DBLβ5 | DBLγ10 | DBLδ1 | CIDRβ1 | | |
| PFKE02.g501 | DBLα0.18 | CIDRα4 | DBLβ5 | DBLδ1 | CIDRβ5 | | | |
| PFKE02.g10 | DBLα0.19 | CIDRα2.8 | DBLδ1 | CIDRβ1 | | | | |
| PFKE02.g14 | DBLα0.2 | CIDRα3.2 | DBLδ6 | CIDRβ2 | | | | |
| PFKE02.g302 | DBLα0.2 | CIDRα3.3 | DBLδ1 | CIDRβ1 | | | | |
| PFKE02.g219 | DBLα0.23 | DBLδ1 | CIDRγ5 | | | | | |
| PFKE02.g87 | DBLα0.3 | CIDRα5 | DBLβ5 | | | | | |
| PFKE02.g195 | DBLα0.3 | CIDRα5 | DBLβ5 | | | | | |
| PFKE02.g256 | DBLα0.3 | CIDRα4 | DBLβ5 | DBLδ5 | CIDRβ4 | | | |
| PFKE02.g570 | DBLα0.3 | CIDRα2.1 | DBLγ2 | DBLγ4 | DBLδ1 | CIDRβ6 | | |
| PFKE02.g5 | DBLα0.4 | CIDRα6 | DBLβ6 | DBLγ12 | DBLδ1 | CIDRγ6 | DBLζ6 | DBLε9 |
| PFKE02.g304 | DBLα0.4 | CIDRα3.1 | DBLδ1 | CIDRβ1 | | | | |
| PFKE02.g353 | DBLα0.4 | CIDRα6 | DBLβ5 | DBLγ13 | DBLδ1 | CIDRβ1 | | |
| PFKE02.g9 | DBLα0.5 | CIDRα2.3 | DBLβ13 | DBLδ1 | CIDRβ7 | | | |
| PFKE02.g260 | DBLα0.5 | CIDRα2.6 | DBLδ1 | CIDRβ6 | | | | |
| PFKE02.g303 | DBLα0.5 | CIDRα2.6 | DBLδ1 | CIDRβ1 | | | | |
| PFKE02.g334 | DBLα0.5 | CIDRα2.5 | DBLδ1 | CIDRγ7 | | | | |
| PFKE02.g476 | DBLα0.5 | CIDRα2.3 | DBLδ1 | CIDRβ1 | | | | |
| PFKE02.g507 | DBLα0.5 | CIDRα2.3 | DBLδ1 | CIDRβ1 | | | | |
| PFKE02.g13 | DBLα0.7 | CIDRα3.1 | DBLδ1 | CIDRβ1 | | | | |
| PFKE02.g15 | DBLα0.8 | CIDRα4 | DBLδ1 | CIDRβ6 | | | | |
| PFKE02.g250 | DBLα0.9 | CIDRα2.1 | DBLδ1 | CIDRγ9 | DBLε12 | | | |
| PFKE02.g569 | DBLα0.9 | CIDRα2.4 | DBLδ1 | CIDRβ1 | | | | |
| PFKE02.g8 | DBLpam1 | DBLpam2 | CIDRpam | DBLpam3 | DBLεpam4 | DBLεpam5 | DBLε10 | |
| PFKE02.g309 | CIDRα2.2 | DBLδ1 | CIDRβ1 | | | | | |
| PFKE02.g37 | CIDRα3.2 | DBLδ1 | CIDRβ6 | | | | | |
| PFKE02.g57 | CIDRα3.2 | DBLδ1 | CIDRβ1 | | | | | |
| PFKE02.g76 | CIDRβ1 | DBLγ10 | | | | | | |
| PFKE02.g36 | CIDRγ1 | DBLζ3 | DBLε6 | | | | | |
| PFKE02.g2 | DBLβ5 | DBLβ5 | DBLβ9 | DBLδ1 | CIDRβ3 | | | |
| PFKE02.g25 | DBLβ5 | DBLγ14 | DBLζ5 | DBLε4 | | | | |
| PFKE02.g149 | DBLβ5 | DBLδ1 | CIDRβ6 | | | | | |
| PFKE02.g310 | DBLβ6 | DBLβ5 | DBLδ1 | CIDRβ6 | | | | |
| PFKE02.g301 | DBLδ1 | CIDRβ6 | | | | | | |
| PFKE02.g291 | DBLδ9 | CIDRα3.2 | DBLδ1 | CIDRβ1 | | | | |
| PFKE03.g10 | DBLα1.2 | CIDRα1.5 | DBLβ3 | DBLγ11 | DBLγ4 | DBLδ6 | | |
| PFKE03.g35 | DBLα1.2 | DBLζ4 | | | | | | |
| PFKE03.g273 | DBLα1.2 | CIDRα1.5 | DBLβ9 | DBLγ11 | DBLγ11 | DBLζ1 | DBLε6 | |
| PFKE03.g170 | DBLα1.3 | DBLε8 | | | | | | |
| PFKE03.g6 | DBLα1.4 | CIDRα1.3 | DBLβ1 | DBLγ15 | DBLε1 | DBLγ8 | DBLζ2 | DBLε5 |
| PFKE03.g9 | DBLα1.4 | CIDRα1.7 | DBLβ1 | DBLβ9 | DBLγ2 | DBLδ1 | CIDRβ3 | |
| PFKE03.g3 | DBLα1.6 | CIDRγ3 | DBLγ7 | DBLδ5 | CIDRβ4 | DBLβ6 | DBLε13 | |
| PFKE03.g323 | DBLα1.6 | CIDRδ1 | DBLβ3 | DBLγ13 | DBLδ1 | CIDRγ7 | | |
| PFKE03.g469 | DBLα1.7 | CIDRα1.4 | DBLβ1 | DBLβ7 | DBLγ11 | DBLδ1 | CIDRβ6 | |
| PFKE03.g274 | DBLα2 | CIDRα1.1 | DBLβ12 | DBLγ4 | DBLδ1 | CIDRβ6 | | |
| PFKE03.g327 | DBLα2 | CIDRα1.1 | DBLβ12 | DBLγ2 | DBLδ1 | CIDRβ1 | | |
| PFKE03.g328 | CIDRβ1 | DBLα0.1 | CIDRα3.1 | DBLδ1 | CIDRβ1 | | | |
| PFKE03.g12 | DBLα0.1 | CIDRα3.1 | DBLδ1 | CIDRβ1 | | | | |
| PFKE03.g63 | DBLα0.1 | CIDRα3.2 | | | | | | |
| PFKE03.g177 | DBLα0.1 | CIDRα3.2 | DBLδ1 | CIDRβ5 | | | | |
| PFKE03.g278 | DBLα0.1 | CIDRα3.1 | DBLε13 | | | | | |
| PFKE03.g286 | DBLα0.1 | CIDRα3.2 | DBLδ1 | CIDRβ1 | | | | |
| PFKE03.g326 | DBLα0.1 | CIDRα3.2 | DBLδ1 | CIDRγ4 | | | | |
| PFKE03.g330 | DBLα0.1 | CIDRα3.1 | DBLδ1 | CIDRβ1 | | | | |
| PFKE03.g515 | DBLα0.1 | CIDRα3.2 | | | | | | |
| PFKE03.g569 | DBLα0.1 | CIDRα3.1 | DBLδ1 | CIDRγ4 | | | | |
| PFKE03.g14 | DBLα0.11 | CIDRα2.4 | DBLδ1 | CIDRβ1 | | | | |
| PFKE03.g17 | DBLα0.11 | CIDRα2.4 | DBLδ1 | CIDRβ1 | | | | |

**Rosetting-associated head structure**

DBLα1.5 /6 /8 — CIDRβ /γ /δ

| Gene | | | | | | | | | |
|---|---|---|---|---|---|---|---|---|---|
| PFKE03.g20 | DBLα0.11 | CIDRα2.4 | DBLδ1 | CIDRβ1 | | | | | |
| PFKE03.g277 | DBLα0.11 | CIDRα2.5 | DBLδ1 | CIDRβ1 | | | | | |
| PFKE03.g280 | DBLα0.11 | CIDRα2.4 | DBLδ1 | CIDRβ1 | | | | | |
| PFKE03.g344 | DBLα0.11 | CIDRα2.4 | DBLδ1 | CIDRγ1 | | | | | |
| PFKE03.g15 | DBLα0.12 | CIDRα2.7 | DBLδ1 | CIDRγ5 | | | | | |
| PFKE03.g16 | DBLα0.12 | CIDRα2.4 | DBLδ1 | CIDRβ1 | | | | | |
| PFKE03.g18 | DBLα0.16 | CIDRα3.4 | DBLδ1 | CIDRβ1 | | | | | |
| PFKE03.g21 | DBLα0.16 | CIDRα3.4 | DBLδ1 | CIDRβ1 | | | | | |
| PFKE03.g325 | DBLα0.16 | CIDRα3.4 | DBLδ1 | CIDRγ5 | | | | | |
| PFKE03.g332 | DBLα0.16 | CIDRα3.4 | DBLδ1 | CIDRβ1 | | | | | |
| PFKE03.g275 | DBLα0.17 | CIDRα4 | DBLδ1 | CIDRβ1 | | | | | |
| PFKE03.g333 | DBLα0.17 | CIDRα3.1 | DBLδ1 | CIDRβ5 | | | | | |
| PFKE03.g381 | DBLα0.17 | CIDRα3.1 | DBLδ1 | CIDRγ5 | | | | | |
| PFKE03.g13 | DBLα0.18 | CIDRα5 | DBLβ5 | DBLγ10 | DBLδ1 | CIDRγ5 | | | |
| PFKE03.g272 | DBLα0.18 | CIDRα5 | DBLβ5 | DBLδ4 | CIDRγ6 | DBLγ11 | DBLζ4 | | |
| PFKE03.g309 | DBLα0.2 | CIDRα3.1 | DBLδ1 | CIDRγ1 | | | | | |
| PFKE03.g329 | DBLα0.2 | CIDRα3.2 | | | | | | | |
| PFKE03.g7 | DBLα0.3 | CIDRα6 | DBLβ4 | DBLγ14 | DBLζ5 | DBLε4 | | | |
| PFKE03.g5 | DBLα0.4 | CIDRα6 | DBLβ5 | DBLγ10 | DBLδ6 | CIDRβ2 | | | |
| PFKE03.g19 | DBLα0.4 | CIDRα4 | DBLβ10 | DBLγ10 | | | | | |
| PFKE03.g118 | DBLα0.5 | CIDRα2.6 | DBLδ1 | CIDRγ5 | | | | | |
| PFKE03.g276 | DBLα0.5 | CIDRα2.9 | DBLδ1 | CIDRβ1 | | | | | |
| PFKE03.g324 | DBLα0.5 | CIDRα2.3 | DBLβ4 | DBLδ1 | CIDRγ7 | DBLε13 | DBLγ12 | | |
| PFKE03.g322 | DBLα0.6 | CIDRα3.2 | DBLδ1 | CIDRγ2 | | | | | |
| PFKE03.g331 | DBLα0.6 | CIDRα3.1 | DBLδ1 | CIDRβ5 | | | | | |
| PFKE03.g8 | DBLα0.8 | CIDRα5 | DBLβ5 | DBLγ13 | DBLδ4 | CIDRγ6 | DBLε2 | DBLζ3 | DBLε9 |
| PFKE03.g281 | DBLα0.9 | CIDRα2.4 | | | | | | | |
| PFKE03.g2 | DBLpam1 | DBLpam2 | CIDRpam | DBLpam3 | DBLεpam4 | DBLεpam5 | DBLε10 | DBLε8 | |
| PFKE03.g11 | DBLpam1 | DBLpam2 | CIDRpam | DBLpam3 | | | | | |
| PFKE03.g377 | DBLpam3 | DBLεpam4 | DBLεpam5 | DBLε10 | | | | | |
| PFKE03.g43 | CIDRα2.4 | DBLδ1 | CIDRβ1 | | | | | | |
| PFKE03.g67 | CIDRα2.4 | DBLδ1 | CIDRγ7 | | | | | | |
| PFKE03.g342 | CIDRα3.2 | DBLδ1 | CIDRβ1 | | | | | | |
| PFKE03.g242 | DBLδ1 | CIDRβ6 | DBLγ10 | | | | | | |
| PFKE03.g334 | DBLδ1 | CIDRγ12 | DBLζ6 | DBLε9 | | | | | |
| PFKE03.g341 | DBLδ1 | CIDRβ1 | | | | | | | |
| PFKE03.g351 | DBLδ1 | CIDRβ1 | | | | | | | |
| PFKE04.g406 | CIDRβ5 | DBLα1.7 | CIDRα1.4 | DBLβ1 | DBLβ6 | DBLβ9 | DBLδ1 | CIDRβ5 | |
| PFKE04.g5 | DBLα1.1 | CIDRα1.6 | DBLβ3 | DBLγ13 | DBLζ3 | DBLε12 | | | |
| PFKE04.g873 | DBLα1.1 | CIDRα1.7 | DBLβ3 | DBLγ2 | DBLγ4 | DBLγ11 | DBLδ1 | CIDRβ1 | |
| PFKE04.g1156 | DBLα1.1 | CIDRα1.4 | DBLβ1 | DBLγ16 | | | | | |
| PFKE04.g1 | DBLα1.2 | CIDRα1.5 | DBLβ6 | DBLγ2 | DBLγ4 | DBLγ11 | DBLδ1 | CIDRβ1 | |
| PFKE04.g3 | DBLα1.2 | CIDRα1.5 | DBLβ6 | DBLγ12 | DBLδ5 | CIDRβ4 | DBLβ7 | DBLγ2 | |
| PFKE04.g845 | DBLα1.3 | DBLε8 | | | | | | | |
| PFKE04.g851 | DBLα1.4 | CIDRα1.3 | DBLβ1 | DBLγ15 | DBLε1 | DBLγ8 | DBLζ1 | DBLε5 | |
| PFKE04.g13 | DBLα1.5 | CIDRγ3 | DBLγ11 | DBLδ5 | CIDRβ4 | | | | |
| PFKE04.g843 | DBLα1.6 | CIDRγ3 | DBLγ7 | DBLε11 | DBLα1.3 | DBLε8 | | | |
| PFKE04.g22 | DBLα1.7 | CIDRα1.1 | DBLβ1 | | | | | | |
| PFKE04.g605 | DBLα1.7 | CIDRα1.4 | DBLβ1 | DBLγ2 | DBLγ6 | DBLγ11 | | | |
| PFKE04.g6 | DBLα1.8 | CIDRγ3 | DBLγ12 | DBLδ5 | CIDRβ4 | DBLβ7 | | | |
| PFKE04.g1042 | DBLα1.8 | CIDRγ3 | DBLγ11 | DBLδ4 | CIDRγ6 | DBLζ6 | DBLε9 | | |
| PFKE04.g597 | DBLα2 | CIDRα1.1 | DBLβ12 | DBLγ6 | DBLδ1 | CIDRδ1 | | | |
| PFKE04.g599 | DBLα2 | CIDRα1.8 | DBLβ12 | DBLδ1 | CIDRβ5 | | | | |
| PFKE04.g600 | DBLα2 | CIDRα1.1 | DBLβ3 | DBLγ11 | DBLζ5 | DBLε4 | | | |
| PFKE04.g302 | DBLα0.1 | CIDRα3.1 | | | | | | | |
| PFKE04.g422 | DBLα0.1 | CIDRα3.2 | DBLδ1 | CIDRγ5 | | | | | |
| PFKE04.g442 | DBLα0.1 | CIDRα3.2 | DBLδ1 | CIDRβ1 | | | | | |
| PFKE04.g614 | DBLα0.1 | CIDRα3.3 | DBLδ1 | CIDRβ1 | | | | | |
| PFKE04.g777 | DBLα0.1 | CIDRα3.1 | DBLδ1 | CIDRβ1 | | | | | |
| PFKE04.g1144 | DBLα0.1 | CIDRα3.3 | DBLδ1 | CIDRβ1 | | | | | |
| PFKE04.g1161 | DBLα0.1 | CIDRα2.2 | DBLε2 | DBLζ3 | DBLε3 | | | | |
| PFKE04.g12 | DBLα0.11 | CIDRα2.4 | DBLδ1 | CIDRβ5 | | | | | |
| PFKE04.g147 | DBLα0.11 | CIDRα2.4 | DBLδ1 | CIDRγ7 | | | | | |
| PFKE04.g598 | DBLα0.12 | CIDRα2.2 | DBLδ1 | CIDRγ1 | | | | | |
| PFKE04.g899 | DBLα0.12 | CIDRα2.2 | DBLδ1 | CIDRγ2 | DBLζ6 | DBLε9 | | | |
| PFKE04.g1155 | DBLα0.12 | CIDRα2.1 | DBLδ1 | CIDRβ3 | | | | | |
| PFKE04.g400 | DBLα0.13 | CIDRα2.9 | DBLδ1 | CIDRβ1 | | | | | |
| PFKE04.g607 | DBLα0.13 | CIDRα2.3 | DBLδ1 | CIDRβ6 | | | | | |
| PFKE04.g602 | DBLα0.14 | CIDRα6 | DBLβ5 | DBLδ1 | CIDRβ1 | | | | |
| PFKE04.g421 | DBLα0.15 | CIDRα3.2 | DBLδ1 | CIDRγ12 | | | | | |
| PFKE04.g16 | DBLα0.16 | CIDRα3.4 | DBLδ1 | CIDRβ1 | DBLα0.9 | CIDRα2.2 | DBLδ1 | CIDRγ5 | |

**PFKE03 (8383)**

**PFKE04 (10668)**

**Rosetting-associated head structure**

DBLα1.5 /6 /8 | CIDRβ /γ /δ

| | | | | | | | | | | | | |
|---|---|---|---|---|---|---|---|---|---|---|---|---|
| PFKE04.g852 | DBLα0.16 | CIDRα3.4 | DBLδ1 | CIDRβ1 | | | | | | | | |
| PFKE04.g8 | DBLα0.17 | CIDRα3.2 | DBLδ1 | CIDRβ1 | | | | | | | | |
| PFKE04.g784 | DBLα0.17 | CIDRα3.1 | DBLδ1 | CIDRβ1 | | | | | | | | |
| PFKE04.g870 | DBLα0.18 | CIDRα5 | DBLβ5 | DBLγ10 | | | | | | | | |
| PFKE04.g1138 | DBLα0.18 | CIDRα4 | DBLδ1 | CIDRγ1 | | | | | | | | |
| PFKE04.g1146 | DBLα0.18 | CIDRα5 | DBLβ5 | DBLγ10 | DBLδ6 | CIDRβ2 | | | | | | |
| PFKE04.g780 | DBLα0.19 | CIDRα2.3 | DBLδ1 | CIDRβ1 | | | | | | | | |
| PFKE04.g4 | DBLα0.22 | CIDRα3.1 | DBLδ1 | CIDRβ1 | | | | | | | | |
| PFKE04.g603 | DBLα0.3 | CIDRα3.1 | DBLδ1 | CIDRγ1 | | | | | | | | |
| PFKE04.g17 | DBLα0.4 | CIDRα6 | DBLβ4 | DBLγ14 | DBLζ3 | DBLε12 | | | | | | |
| PFKE04.g219 | DBLα0.4 | CIDRα3.1 | DBLδ1 | CIDRβ5 | DBLα0.9 | CIDRα5 | DBLβ5 | DBLδ9 | CIDRγ2 | DBLγ13 | DBLζ1 | DBLε9 |
| PFKE04.g604 | DBLα0.4 | CIDRα6 | DBLδ1 | CIDRγ12 | DBLγ3 | DBLζ4 | | | | | | |
| PFKE04.g647 | DBLα0.4 | CIDRα5 | DBLβ5 | | | | | | | | | |
| PFKE04.g7 | DBLα0.5 | CIDRα2.6 | DBLγ11 | DBLδ1 | CIDRβ1 | | | | | | | |
| PFKE04.g10 | DBLα0.5 | CIDRα2.3 | DBLδ1 | CIDRγ4 | | | | | | | | |
| PFKE04.g606 | DBLα0.5 | CIDRα2.3 | DBLδ1 | CIDRγ9 | | | | | | | | |
| PFKE04.g775 | DBLα0.5 | CIDRα2.6 | DBLβ13 | DBLδ1 | CIDRγ10 | | | | | | | |
| PFKE04.g779 | DBLα0.5 | CIDRα2.3 | DBLδ1 | CIDRβ6 | | | | | | | | |
| PFKE04.g366 | DBLα0.6 | CIDRα3.1 | DBLδ1 | CIDRβ1 | | | | | | | | |
| PFKE04.g1176 | DBLα0.7 | CIDRα4 | DBLδ1 | CIDRβ3 | | | | | | | | |
| PFKE04.g887 | DBLα0.8 | CIDRα3.4 | DBLδ1 | CIDRβ1 | | | | | | | | |
| PFKE04.g21 | DBLα0.9 | CIDRα3.2 | DBLδ1 | CIDRβ1 | | | | | | | | |
| PFKE04.g428 | DBLα0.9 | CIDRα5 | | | | | | | | | | |
| PFKE04.g499 | DBLα0.9 | CIDRα2.7 | DBLδ1 | CIDRβ1 | | | | | | | | |
| PFKE04.g601 | DBLα0.9 | CIDRα2.3 | DBLδ1 | CIDRβ1 | | | | | | | | |
| PFKE04.g610 | DBLα0.9 | CIDRα2.2 | DBLδ1 | CIDRβ1 | | | | | | | | |
| PFKE04.g613 | DBLα0.9 | CIDRα2.1 | DBLδ1 | CIDRγ10 | | | | | | | | |
| PFKE04.g776 | DBLα0.9 | CIDRα2.7 | DBLγ14 | DBLζ5 | DBLε4 | | | | | | | |
| PFKE04.g940 | DBLα0.9 | CIDRα2.3 | DBLδ1 | CIDRβ5 | | | | | | | | |
| PFKE04.g1171 | DBLpam1 | DBLpam2 | CIDRpam | DBLpam3 | DBLεpam4 | DBLεpam5 | DBLε10 | | | | | |
| PFKE04.g2 | CIDRα1.7 | DBLβ1 | DBLγ11 | DBLγ12 | DBLδ4 | CIDRδ1 | DBLβ9 | DBLγ13 | | | | |
| PFKE04.g424 | DBLβ5 | DBLγ16 | DBLδ1 | CIDRγ4 | | | | | | | | |
| PFKE04.g748 | DBLβ6 | | | | | | | | | | | |
| PFKE04.g11 | DBLβ8 | DBLβ5 | DBLδ1 | CIDRβ1 | | | | | | | | |
| PFKE04.g231 | DBLδ1 | CIDRβ1 | | | | | | | | | | |
| PFKE04.g9 | DBLγ11 | DBLγ11 | DBLδ1 | CIDRγ6 | | | | | | | | |
| PFKE04.g795 | DBLγ11 | DBLδ1 | CIDRβ5 | | | | | | | | | |

| | | | | | | | | | |
|---|---|---|---|---|---|---|---|---|---|
| PFKE05.g2 | DBLα1.4 | CIDRα1.7 | DBLβ1 | DBLβ6 | DBLγ11 | DBLγ2 | DBLδ6 | CIDRβ4 | |
| PFKE05.g185 | DBLα1.4 | CIDRα1.6 | DBLβ3 | DBLγ11 | DBLγ12 | DBLδ5 | CIDRβ3 | DBLβ7 | DBLγ9 |
| PFKE05.g186 | DBLα1.5 | CIDRγ3 | DBLβ7 | DBLγ11 | DBLδ3 | CIDRγ2 | DBLζ6 | DBLε6 | |
| PFKE05.g187 | DBLα1.7 | CIDRα1.4 | DBLβ1 | DBLγ11 | DBLγ2 | DBLδ1 | CIDRβ1 | | |
| PFKE05.g286 | DBLα1.7 | CIDRα1.4 | DBLβ1 | DBLβ7 | DBLβ6 | DBLδ1 | | | |
| PFKE05.g1 | DBLα1.8 | CIDRβ2 | DBLγ7 | DBLε11 | | | | | |
| PFKE05.g355 | DBLα2 | CIDRα1.1 | DBLβ12 | DBLγ6 | DBLδ1 | CIDRβ6 | | | |
| PFKE05.g13 | DBLα0.1 | CIDRα3.1 | DBLε13 | | | | | | |
| PFKE05.g221 | DBLα0.1 | CIDRα3.2 | DBLδ1 | CIDRγ7 | DBLα0.17 | CIDRα3.1 | DBLδ1 | CIDRγ4 | |
| PFKE05.g228 | DBLα0.1 | CIDRα3.1 | DBLδ1 | CIDRβ1 | | | | | |
| PFKE05.g232 | DBLα0.1 | CIDRα2.2 | DBLδ1 | CIDRβ1 | | | | | |
| PFKE05.g3 | DBLα0.13 | CIDRα2.6 | DBLδ1 | CIDRβ1 | | | | | |
| PFKE05.g10 | DBLα0.15 | CIDRα3.2 | DBLδ1 | CIDRβ1 | | | | | |
| PFKE05.g11 | DBLα0.15 | CIDRα3.2 | DBLδ1 | CIDRβ6 | | | | | |
| PFKE05.g195 | DBLα0.16 | CIDRα3.4 | DBLδ1 | CIDRβ1 | | | | | |
| PFKE05.g231 | DBLα0.17 | CIDRα3.2 | DBLδ1 | CIDRβ5 | | | | | |
| PFKE05.g237 | DBLα0.17 | CIDRα3.3 | DBLδ1 | CIDRβ1 | | | | | |
| PFKE05.g357 | DBLα0.17 | CIDRα3.1 | DBLδ1 | CIDRβ1 | | | | | |
| PFKE05.g8 | DBLα0.18 | CIDRα6 | DBLβ3 | DBLγ11 | DBLδ8 | CIDRβ2 | | | |
| PFKE05.g235 | DBLα0.18 | CIDRα4 | DBLβ5 | DBLδ1 | CIDRβ6 | | | | |
| PFKE05.g188 | DBLα0.19 | CIDRα2.3 | DBLδ1 | CIDRγ5 | | | | | |
| PFKE05.g234 | DBLα0.19 | CIDRα2.5 | DBLδ1 | CIDRβ1 | | | | | |
| PFKE05.g226 | DBLα0.2 | CIDRα3.1 | DBLδ1 | CIDRβ1 | | | | | |
| PFKE05.g229 | DBLα0.22 | CIDRα3.3 | DBLδ1 | CIDRβ1 | | | | | |
| PFKE05.g363 | DBLα0.22 | CIDRα3.1 | DBLβ5 | DBLγ10 | DBLδ2 | CIDRγ6 | | | |
| PFKE05.g14 | DBLα0.3 | CIDRα3.2 | DBLγ13 | DBLδ1 | CIDRβ1 | | | | |
| PFKE05.g362 | DBLα0.4 | CIDRα6 | DBLβ5 | DBLγ2 | DBLδ8 | CIDRβ3 | | | |
| PFKE05.g31 | DBLα0.5 | CIDRα2.1 | DBLβ5 | DBLδ1 | CIDRβ1 | | | | |
| PFKE05.g48 | DBLα0.5 | CIDRα2.1 | DBLβ5 | DBLδ1 | | | | | |
| PFKE05.g84 | DBLα0.5 | CIDRα2.1 | DBLβ5 | | | | | | |
| PFKE05.g254 | DBLα0.5 | CIDRα2.5 | DBLβ8 | DBLδ1 | CIDRβ1 | | | | |
| PFKE05.g284 | DBLα0.5 | CIDRα2.5 | DBLβ13 | DBLγ5 | DBLδ1 | CIDRβ1 | | | |
| PFKE05.g292 | DBLα0.5 | CIDRα2.3 | DBLβ8 | DBLγ5 | DBLδ1 | CIDRγ5 | | | |
| PFKE05.g393 | DBLα0.5 | CIDRα2.3 | DBLδ1 | CIDRγ1 | DBLγ3 | DBLζ4 | | | |

**Rosetting-associated head structure**

**DBLα1.5 /6 /8** **CIDRβ /γ /δ**

### PFKE05
# (11014)

| | | | | | | | | | | | | | | | |
|---|---|---|---|---|---|---|---|---|---|---|---|---|---|---|---|
| PFKE05.g7 | DBLα0.6 | CIDRα4 | DBLγ5 | DBLδ1 | CIDRβ1 | | | | | | | | | | |
| PFKE05.g189 | DBLα0.6 | CIDRα3.1 | DBLδ1 | CIDRβ1 | | | | | | | | | | | |
| PFKE05.g227 | DBLα0.6 | CIDRα3.1 | DBLδ1 | CIDRβ1 | | | | | | | | | | | |
| PFKE05.g230 | DBLα0.8 | CIDRα3.4 | DBLβ5 | DBLγ10 | | | | | | | | | | | |
| PFKE05.g173 | DBLα0.9 | CIDRα2.1 | DBLδ1 | CIDRγ1 | | | | | | | | | | | |
| PFKE05.g233 | DBLα0.9 | CIDRα2.2 | DBLδ1 | | | | | | | | | | | | |
| PFKE05.g283 | DBLα0.9 | CIDRα2.7 | DBLβ5 | DBLδ1 | CIDRβ1 | DBLγ3 | DBLζ4 | | | | | | | | |
| PFKE05.g194 | CIDRα1.7 | DBLβ1 | DBLβ9 | DBLδ1 | | | | | | | | | | | |
| PFKE05.g33 | CIDRγ3 | DBLγ15 | DBLε1 | DBLε11 | DBLζ2 | DBLε6 | | | | | | | | | |
| PFKE05.g22 | DBLδ1 | CIDRγ4 | | | | | | | | | | | | | |
| PFKE05.g24 | DBLδ1 | CIDRβ1 | | | | | | | | | | | | | |
| PFKE05.g44 | DBLδ1 | CIDRβ4 | | | | | | | | | | | | | |
| PFKE05.g61 | DBLδ1 | CIDRβ1 | | | | | | | | | | | | | |
| PFKE06.g3 | DBLα1.1 | CIDRα1.4 | DBLβ12 | DBLγ2 | DBLγ12 | DBLδ1 | CIDRβ3 | | | | | | | | |
| PFKE06.g270 | DBLα1.2 | CIDRα1.5 | DBLβ6 | DBLγ2 | DBLγ14 | DBLζ5 | DBLε4 | | | | | | | | |
| PFKE06.g235 | DBLα1.3 | DBLε8 | | | | | | | | | | | | | |
| PFKE06.g2 | DBLα1.4 | CIDRα1.3 | DBLβ1 | DBLγ15 | DBLε1 | DBLγ8 | DBLζ1 | DBLε5 | | | | | | | |
| PFKE06.g335 | DBLα1.5 | CIDRδ1 | DBLγ11 | DBLγ13 | DBLδ1 | CIDRβ1 | | | | | | | | | |
| PFKE06.g5 | DBLα1.6 | CIDRδ1 | DBLγ12 | DBLδ5 | CIDRβ4 | DBLβ9 | DBLγ2 | | | | | | | | |
| PFKE06.g6 | DBLα1.6 | CIDRγ3 | DBLβ6 | DBLδ4 | CIDRγ2 | DBLβ6 | | | | | | | | | |
| PFKE06.g8 | DBLα1.6 | CIDRδ1 | DBLβ6 | DBLγ12 | DBLγ11 | DBLγ11 | | | | | | | | | |
| PFKE06.g409 | DBLα1.6 | CIDRδ1 | DBLγ12 | DBLδ5 | CIDRβ4 | DBLβ7 | DBLγ6 | DBLα1.2 | CIDRα1.6 | DBLγ7 | DBLδ4 | CIDRγ1 | DBLζ2 | DBLε11 | DBLε3 |
| PFKE06.g229 | DBLα1.7 | CIDRα1.4 | DBLβ3 | DBLβ9 | DBLγ11 | DBLγ9 | DBLδ1 | CIDRβ6 | | | | | | | |
| PFKE06.g233 | DBLα1.8 | CIDRβ4 | DBLγ11 | DBLγ2 | DBLγ6 | | | | | | | | | | |
| PFKE06.g334 | DBLα2 | CIDRα1.8 | DBLβ4 | DBLγ3 | DBLγ13 | DBLζ2 | DBLε4 | | | | | | | | |
| PFKE06.g340 | DBLα2 | CIDRα1.1 | DBLβ12 | DBLγ4 | DBLδ1 | CIDRβ1 | | | | | | | | | |
| PFKE06.g20 | DBLα0.1 | CIDRα3.1 | DBLδ1 | CIDRγ1 | | | | | | | | | | | |
| PFKE06.g274 | DBLα0.1 | CIDRα3.2 | DBLδ1 | CIDRβ5 | | | | | | | | | | | |
| PFKE06.g341 | DBLα0.1 | CIDRα3.1 | DBLδ1 | CIDRβ1 | | | | | | | | | | | |
| PFKE06.g552 | DBLα0.1 | CIDRα3.1 | | | | | | | | | | | | | |
| PFKE06.g17 | DBLα0.1 | CIDRα2.4 | DBLδ1 | CIDRβ1 | | | | | | | | | | | |
| PFKE06.g232 | DBLα0.1 | CIDRα2.2 | DBLδ1 | CIDRβ1 | | | | | | | | | | | |
| PFKE06.g230 | DBLα0.11 | CIDRα2.1 | DBLβ8 | DBLγ5 | DBLδ1 | CIDRβ1 | | | | | | | | | |
| PFKE06.g553 | DBLα0.12 | CIDRα2.1 | DBLβ12 | DBLγ12 | DBLδ4 | CIDRδ1 | DBLε2 | DBLε7 | DBLε3 | | | | | | |
| PFKE06.g336 | DBLα0.13 | CIDRα2.3 | DBLβ4 | DBLδ1 | CIDRγ5 | | | | | | | | | | |
| PFKE06.g16 | DBLα0.15 | CIDRα3.2 | DBLδ1 | CIDRβ6 | | | | | | | | | | | |
| PFKE06.g234 | DBLα0.15 | CIDRα3.2 | DBLδ1 | CIDRγ4 | | | | | | | | | | | |
| PFKE06.g272 | DBLα0.15 | CIDRα3.2 | DBLδ1 | CIDRβ1 | | | | | | | | | | | |
| PFKE06.g275 | DBLα0.15 | CIDRα3.2 | DBLδ1 | CIDRβ1 | | | | | | | | | | | |
| PFKE06.g14 | DBLα0.16 | CIDRα3.4 | DBLδ1 | CIDRβ1 | | | | | | | | | | | |
| PFKE06.g276 | DBLα0.16 | CIDRα3.4 | DBLδ1 | CIDRβ1 | | | | | | | | | | | |
| PFKE06.g11 | DBLα0.17 | CIDRα3.2 | DBLδ1 | CIDRγ2 | | | | | | | | | | | |
| PFKE06.g145 | DBLα0.17 | CIDRα3.3 | DBLδ1 | CIDRγ6 | DBLζ6 | DBLε6 | | | | | | | | | |
| PFKE06.g231 | DBLα0.18 | CIDRα5 | DBLβ5 | DBLγ10 | DBLδ1 | CIDRβ1 | | | | | | | | | |
| PFKE06.g267 | DBLα0.18 | CIDRα6 | DBLβ5 | DBLγ12 | DBLδ4 | CIDRγ6 | DBLε2 | DBLε7 | DBLε3 | | | | | | |
| PFKE06.g278 | DBLα0.18 | CIDRα5 | DBLβ5 | DBLγ10 | | | | | | | | | | | |
| PFKE06.g273 | DBLα0.19 | CIDRα6 | DBLδ1 | CIDRβ1 | | | | | | | | | | | |
| PFKE06.g282 | DBLα0.19 | CIDRα2.8 | DBLδ1 | CIDRβ5 | | | | | | | | | | | |
| PFKE06.g18 | DBLα0.2 | CIDRα3.1 | DBLδ1 | CIDRβ3 | | | | | | | | | | | |
| PFKE06.g277 | DBLα0.2 | CIDRα3.3 | DBLδ1 | CIDRβ1 | | | | | | | | | | | |
| PFKE06.g280 | DBLα0.2 | CIDRα4 | DBLδ1 | CIDRβ6 | | | | | | | | | | | |
| PFKE06.g292 | DBLα0.2 | CIDRα3.1 | DBLδ1 | CIDRβ1 | | | | | | | | | | | |
| PFKE06.g9 | DBLα0.21 | CIDRα6 | DBLβ5 | DBLγ12 | DBLδ4 | CIDRβ4 | DBLε2 | | | | | | | | |
| PFKE06.g12 | DBLα0.21 | CIDRα2.1 | DBLβ2 | DBLδ1 | CIDRβ4 | | | | | | | | | | |
| PFKE06.g269 | DBLα0.4 | CIDRα6 | DBLβ5 | DBLγ10 | DBLδ7 | CIDRβ4 | | | | | | | | | |
| PFKE06.g7 | DBLα0.5 | CIDRα2.3 | DBLγ11 | DBLδ1 | CIDRβ1 | | | | | | | | | | |
| PFKE06.g144 | DBLα0.5 | CIDRα2.3 | DBLδ1 | CIDRβ1 | | | | | | | | | | | |
| PFKE06.g281 | DBLα0.5 | CIDRα2.9 | DBLδ1 | CIDRβ1 | | | | | | | | | | | |
| PFKE06.g15 | DBLα0.6 | CIDRα3.1 | DBLδ1 | CIDRβ6 | | | | | | | | | | | |
| PFKE06.g249 | DBLα0.6 | CIDRα3.2 | DBLδ1 | CIDRβ1 | | | | | | | | | | | |
| PFKE06.g238 | DBLα0.8 | CIDRα2.5 | DBLδ1 | CIDRγ5 | | | | | | | | | | | |
| PFKE06.g10 | DBLα0.9 | CIDRα2.7 | DBLδ1 | CIDRβ1 | | | | | | | | | | | |
| PFKE06.g34 | DBLα0.9 | CIDRα4 | DBLδ1 | CIDRγ5 | | | | | | | | | | | |
| PFKE06.g4 | DBLpam1 | DBLpam2 | CIDRpam | DBLpam3 | DBLεpam4 | DBLεpam5 | DBLε10 | | | | | | | | |
| PFKE06.g114 | CIDRα2.2 | DBLδ3 | | | | | | | | | | | | | |
| PFKE06.g115 | CIDRα2.2 | DBLδ1 | | | | | | | | | | | | | |
| PFKE06.g251 | CIDRα2.5 | DBLδ1 | CIDRβ3 | | | | | | | | | | | | |
| PFKE06.g42 | CIDRγ9 | DBLζ6 | DBLε9 | | | | | | | | | | | | |
| PFKE06.g41 | DBLβ5 | DBLβ5 | DBLδ1 | CIDRβ2 | | | | | | | | | | | |
| PFKE06.g65 | DBLβ5 | DBLβ5 | DBLδ1 | CIDRβ1 | | | | | | | | | | | |
| PFKE06.g271 | DBLβ5 | DBLβ5 | DBLδ1 | CIDRβ1 | | | | | | | | | | | |

**Rosetting-associated head structure**

DBLα1.5 /6 /8 CIDRβ /γ /δ

### PFKE06 (10936)

| | | | | | | | | | |
|---|---|---|---|---|---|---|---|---|---|
| PFKE06.g339 | DBLβ5 | DBLδ1 | CIDRβ1 | | | | | | |
| PFKE06.g61 | DBLδ1 | CIDRβ1 | | | | | | | |
| PFKE06.g239 | DBLδ1 | CIDRβ1 | | | | | | | |
| PFKE06.g314 | DBLδ1 | CIDRβ1 | | | | | | | |
| PFKE07.g60 | DBLα1.1 | CIDRα1.2 | DBLβ11 | DBLγ1 | DBLε1 | DBLγ8 | DBLζ2 | DBLε5 | |
| PFKE07.g255 | DBLα1.1 | CIDRα1.7 | DBLβ1 | DBLγ16 | DBLδ5 | CIDRβ4 | DBLβ7 | DBLγ5 | |
| PFKE07.g322 | DBLα1.1 | CIDRα1.7 | DBLζ3 | DBLγ2 | DBLγ4 | DBLγ2 | DBLδ1 | CIDRβ1 | |
| PFKE07.g790 | DBLα1.1 | CIDRα1.7 | DBLβ1 | DBLγ13 | DBLδ3 | CIDRγ2 | | | |
| PFKE07.g853 | DBLα1.1 | CIDRα1.4 | DBLγ12 | DBLδ5 | CIDRβ3 | DBLβ9 | | | |
| PFKE07.g297 | DBLα1.2 | CIDRα1.6 | DBLβ3 | DBLγ6 | DBLδ5 | CIDRβ3 | DBLβ6 | DBLγ11 | DBLε4 |
| PFKE07.g407 | DBLα1.2 | CIDRα1.5 | DBLβ6 | DBLγ16 | DBLδ5 | CIDRβ3 | DBLβ7 | DBLγ2 | |
| PFKE07.g550 | DBLα1.2 | CIDRα1.5 | DBLβ6 | DBLγ2 | DBLγ4 | DBLδ1 | CIDRβ1 | | |
| PFKE07.g641 | DBLα1.2 | CIDRα1.7 | DBLζ3 | DBLγ2 | DBLγ4 | DBLδ1 | CIDRγ5 | | |
| PFKE07.g217 | DBLα1.3 | DBLε8 | | | | | | | |
| PFKE07.g216 | DBLα1.4 | CIDRα1.4 | DBLβ3 | | | | | | |
| PFKE07.g569 | DBLα1.4 | CIDRα1.7 | DBLβ1 | DBLβ6 | DBLγ11 | DBLδ1 | CIDRβ1 | | |
| PFKE07.g1073 | DBLα1.4 | CIDRα1.7 | DBLβ1 | DBLγ11 | DBLγ12 | DBLδ4 | CIDRδ1 | DBLβ9 | DBLγ13 |
| PFKE07.g692 | DBLα1.5 | CIDRγ3 | DBLγ12 | DBLδ5 | CIDRβ4 | DBLβ9 | DBLγ2 | | |
| PFKE07.g874 | DBLα1.5 | CIDRδ1 | DBLγ11 | DBLγ2 | DBLδ1 | CIDRβ5 | | | |
| PFKE07.g367 | DBLα1.6 | CIDRγ3 | DBLβ6 | DBLδ4 | CIDRγ6 | DBLβ6 | | | |
| PFKE07.g1043 | DBLα1.6 | CIDRγ3 | DBLβ7 | DBLγ13 | | | | | |
| PFKE07.g1048 | DBLα1.6 | CIDRδ2 | DBLγ2 | DBLδ1 | CIDRβ1 | | | | |
| PFKE07.g597 | DBLα1.7 | CIDRα1.4 | DBLβ1 | DBLγ2 | DBLγ10 | DBLδ1 | CIDRβ1 | | |
| PFKE07.g613 | DBLα1.7 | CIDRα1.4 | DBLβ1 | DBLβ3 | DBLγ11 | DBLδ1 | CIDRβ5 | | |
| PFKE07.g615 | DBLα1.7 | CIDRα1.4 | DBLβ1 | DBLβ6 | DBLγ11 | DBLγ11 | DBLδ1 | CIDRβ1 | |
| PFKE07.g26 | DBLα1.8 | CIDRβ2 | DBLγ7 | DBLε11 | DBLζ2 | DBLε6 | | | |
| PFKE07.g667 | DBLα1.8 | CIDRβ2 | DBLγ7 | DBLε11 | DBLζ3 | DBLε6 | | | |
| PFKE07.g73 | DBLα2 | CIDRα1.1 | DBLβ1 | DBLγ4 | DBLδ1 | CIDRβ1 | | | |
| PFKE07.g901 | DBLα2 | CIDRα1.1 | DBLβ12 | DBLγ4 | DBLδ1 | CIDRβ1 | | | |
| PFKE07.g33 | DBLα0.1 | CIDRα3.3 | DBLδ1 | CIDRβ1 | | | | | |
| PFKE07.g52 | DBLα0.1 | CIDRα3.1 | DBLδ1 | CIDRγ10 | | | | | |
| PFKE07.g57 | DBLα0.1 | CIDRα3.2 | DBLδ1 | CIDRγ4 | | | | | |
| PFKE07.g81 | DBLα0.1 | CIDRα3.1 | DBLδ1 | CIDRγ4 | | | | | |
| PFKE07.g201 | DBLα0.1 | CIDRα3.1 | DBLδ1 | | | | | | |
| PFKE07.g295 | DBLα0.1 | CIDRα3.2 | DBLδ1 | | | | | | |
| PFKE07.g368 | DBLα0.1 | CIDRα3.1 | DBLδ1 | CIDRβ1 | | | | | |
| PFKE07.g469 | DBLα0.1 | CIDRα4 | | | | | | | |
| PFKE07.g996 | DBLα0.1 | CIDRα3.2 | DBLβ5 | DBLδ1 | CIDRβ6 | | | | |
| PFKE07.g72 | DBLα0.1 | CIDRα2.2 | DBLδ1 | CIDRβ1 | | | | | |
| PFKE07.g95 | DBLα0.1 | CIDRα2.2 | DBLδ1 | CIDRβ1 | | | | | |
| PFKE07.g48 | DBLα0.11 | CIDRα2.4 | DBLδ1 | CIDRγ10 | | | | | |
| PFKE07.g58 | DBLα0.11 | CIDRα2.4 | DBLδ1 | CIDRβ6 | | | | | |
| PFKE07.g843 | DBLα0.11 | CIDRα2.8 | DBLδ1 | CIDRβ5 | | | | | |
| PFKE07.g1034 | DBLα0.11 | CIDRα2.4 | DBLβ5 | DBLδ1 | CIDRβ1 | | | | |
| PFKE07.g41 | DBLα0.12 | CIDRα2.2 | DBLδ1 | CIDRβ1 | | | | | |
| PFKE07.g50 | DBLα0.12 | CIDRα2.1 | DBLδ1 | CIDRγ10 | | | | | |
| PFKE07.g68 | DBLα0.12 | CIDRα2.1 | DBLδ1 | CIDRβ5 | | | | | |
| PFKE07.g1072 | DBLα0.12 | CIDRα2.1 | DBLδ1 | CIDRβ1 | | | | | |
| PFKE07.g49 | DBLα0.13 | CIDRα2.3 | DBLδ1 | CIDRβ6 | | | | | |
| PFKE07.g40 | DBLα0.14 | CIDRα3.2 | DBLδ1 | CIDRβ1 | | | | | |
| PFKE07.g69 | DBLα0.14 | CIDRα4 | DBLδ1 | CIDRγ4 | | | | | |
| PFKE07.g12 | DBLα0.15 | CIDRα3.2 | DBLδ1 | CIDRβ1 | | | | | |
| PFKE07.g43 | DBLα0.15 | CIDRα3.2 | DBLδ1 | CIDRβ1 | | | | | |
| PFKE07.g70 | DBLα0.15 | CIDRα3.2 | DBLδ1 | CIDRγ5 | | | | | |
| PFKE07.g71 | DBLα0.15 | CIDRα2.5 | DBLδ1 | CIDRβ5 | | | | | |
| PFKE07.g77 | DBLα0.15 | CIDRα3.2 | DBLδ1 | CIDRβ1 | | | | | |
| PFKE07.g932 | DBLα0.15 | CIDRα3.1 | DBLδ1 | CIDRγ12 | | | | | |
| PFKE07.g13 | DBLα0.16 | CIDRα3.4 | DBLδ1 | CIDRβ3 | | | | | |
| PFKE07.g61 | DBLα0.16 | CIDRα3.4 | DBLδ1 | CIDRβ1 | | | | | |
| PFKE07.g940 | DBLα0.16 | CIDRα5 | DBLβ5 | DBLγ11 | DBLδ1 | CIDRβ1 | | | |
| PFKE07.g1070 | DBLα0.16 | CIDRα3.4 | DBLδ1 | CIDRγ7 | | | | | |
| PFKE07.g39 | DBLα0.17 | CIDRα3.3 | DBLδ1 | CIDRβ1 | | | | | |
| PFKE07.g63 | DBLα0.17 | CIDRα3.3 | DBLδ1 | CIDRβ1 | | | | | |
| PFKE07.g470 | DBLα0.17 | CIDRα3.1 | DBLδ1 | CIDRβ1 | | | | | |
| PFKE07.g1049 | DBLα0.17 | CIDRα3.1 | DBLδ1 | CIDRγ12 | | | | | |
| PFKE07.g1067 | DBLα0.17 | CIDRα3.1 | DBLδ1 | CIDRβ3 | | | | | |
| PFKE07.g1071 | DBLα0.17 | CIDRα3.3 | DBLδ1 | CIDRβ1 | | | | | |
| PFKE07.g75 | DBLα0.18 | CIDRα3.1 | DBLδ1 | CIDRβ1 | | | | | |
| PFKE07.g85 | DBLα0.18 | CIDRα5 | DBLβ6 | DBLγ9 | | | | | |
| PFKE07.g1035 | DBLα0.18 | CIDRα6 | DBLβ5 | DBLδ1 | CIDRγ7 | | | | |
| PFKE07.g1051 | DBLα0.18 | CIDRα4 | DBLβ5 | DBLδ1 | CIDRβ1 | | | | |

**Rosetting-associated head structure**

**DBLα1.5 /6 /8** — **CIDRβ /γ /δ**

**PFKE07 (10975)**

| | | | | | | | | | | | | | | | | |
|---|---|---|---|---|---|---|---|---|---|---|---|---|---|---|---|---|
| PFKE07.g1057 | DBLα0.18 | CIDRα6 | DBLβ5 | DBLδ1 | CIDRβ5 | | | | | | | | | | | |
| PFKE07.g55 | DBLα0.19 | CIDRα2.5 | DBLδ1 | CIDRβ3 | | | | | | | | | | | | |
| PFKE07.g779 | DBLα0.19 | CIDRα2.9 | DBLδ1 | CIDRβ1 | DBLγ3 | DBLζ4 | | | | | | | | | | |
| PFKE07.g62 | DBLα0.2 | CIDRα3.1 | DBLδ1 | CIDRβ1 | | | | | | | | | | | | |
| PFKE07.g1068 | DBLα0.2 | CIDRα3.3 | DBLδ1 | CIDRβ1 | | | | | | | | | | | | |
| PFKE07.g202 | DBLα0.22 | CIDRα3.1 | DBLβ5 | DBLδ5 | CIDRβ4 | DBLγ3 | DBLζ4 | | | | | | | | | |
| PFKE07.g260 | DBLα0.22 | CIDRα3.1 | DBLδ1 | CIDRβ1 | | | | | | | | | | | | |
| PFKE07.g750 | DBLα0.3 | CIDRα2.1 | DBLβ5 | DBLδ1 | CIDRγ9 | DBLε2 | DBLε7 | DBLε3 | | | | | | | | |
| PFKE07.g800 | DBLα0.3 | CIDRα5 | DBLβ5 | DBLγ2 | DBLδ8 | CIDRβ4 | | | | | | | | | | |
| PFKE07.g51 | DBLα0.4 | CIDRα6 | DBLδ1 | CIDRβ1 | | | | | | | | | | | | |
| PFKE07.g56 | DBLα0.4 | CIDRα3.2 | DBLδ1 | CIDRβ3 | | | | | | | | | | | | |
| PFKE07.g78 | DBLα0.4 | CIDRα6 | DBLδ1 | CIDRβ1 | | | | | | | | | | | | |
| PFKE07.g118 | DBLα0.4 | CIDRα5 | DBLβ3 | DBLγ11 | | | | | | | | | | | | |
| PFKE07.g121 | DBLα0.4 | CIDRα6 | DBLδ1 | CIDRβ6 | | | | | | | | | | | | |
| PFKE07.g955 | DBLα0.4 | CIDRα6 | DBLβ5 | DBLγ2 | DBLδ8 | CIDRβ2 | | | | | | | | | | |
| PFKE07.g985 | DBLα0.4 | CIDRα5 | DBLβ6 | DBLγ9 | | | | | | | | | | | | |
| PFKE07.g1066 | DBLα0.4 | CIDRα6 | DBLγ13 | DBLζ2 | DBLε4 | | | | | | | | | | | |
| PFKE07.g20 | DBLα0.5 | CIDRα2.5 | DBLδ1 | CIDRβ1 | | | | | | | | | | | | |
| PFKE07.g30 | DBLα0.5 | CIDRα2.6 | DBLδ1 | CIDRβ1 | | | | | | | | | | | | |
| PFKE07.g42 | DBLα0.5 | CIDRα2.3 | DBLδ1 | CIDRβ1 | | | | | | | | | | | | |
| PFKE07.g64 | DBLα0.5 | CIDRα2.6 | DBLδ1 | CIDRγ5 | | | | | | | | | | | | |
| PFKE07.g96 | DBLα0.5 | CIDRα2.1 | DBLδ1 | CIDRγ1 | | | | | | | | | | | | |
| PFKE07.g1002 | DBLα0.5 | CIDRα2.3 | DBLβ13 | DBLδ1 | CIDRβ1 | | | | | | | | | | | |
| PFKE07.g1023 | DBLα0.5 | CIDRα2.3 | DBLβ8 | DBLδ1 | CIDRβ1 | | | | | | | | | | | |
| PFKE07.g1064 | DBLα0.5 | CIDRα2.5 | DBLδ1 | CIDRβ1 | | | | | | | | | | | | |
| PFKE07.g226 | DBLα0.6 | CIDRα3.2 | DBLβ5 | DBLγ10 | DBLδ4 | CIDRγ1 | DBLε2 | DBLζ3 | DBLε9 | | | | | | | |
| PFKE07.g424 | DBLα0.6 | CIDRα3.1 | DBLβ4 | DBLγ13 | DBLδ1 | CIDRβ1 | | | | | | | | | | |
| PFKE07.g751 | DBLα0.6 | CIDRα3.2 | DBLδ1 | CIDRβ1 | | | | | | | | | | | | |
| PFKE07.g1069 | DBLα0.7 | CIDRα3.4 | DBLδ1 | CIDRβ3 | | | | | | | | | | | | |
| PFKE07.g84 | DBLα0.8 | CIDRα4 | DBLδ1 | CIDRβ1 | | | | | | | | | | | | |
| PFKE07.g94 | DBLα0.8 | CIDRα4 | DBLδ1 | CIDRβ5 | | | | | | | | | | | | |
| PFKE07.g170 | DBLα0.8 | DBLδ1 | CIDRβ1 | | | | | | | | | | | | | |
| PFKE07.g1058 | DBLα0.8 | CIDRα3.5 | DBLδ1 | CIDRβ1 | | | | | | | | | | | | |
| PFKE07.g38 | DBLα0.9 | CIDRα2.3 | DBLδ1 | CIDRγ12 | | | | | | | | | | | | |
| PFKE07.g76 | DBLα0.9 | CIDRα2.2 | DBLδ1 | CIDRβ5 | | | | | | | | | | | | |
| PFKE07.g82 | DBLα0.9 | CIDRα2.3 | DBLδ1 | CIDRβ1 | | | | | | | | | | | | |
| PFKE07.g83 | DBLα0.9 | CIDRα2.2 | DBLδ1 | CIDRγ1 | | | | | | | | | | | | |
| PFKE07.g222 | DBLα0.9 | CIDRα2.2 | DBLδ1 | | | | | | | | | | | | | |
| PFKE07.g296 | DBLα0.9 | CIDRα2.2 | | | | | | | | | | | | | | |
| PFKE07.g1007 | DBLα0.9 | CIDRα2.2 | DBLδ1 | CIDRγ5 | | | | | | | | | | | | |
| PFKE07.g1050 | DBLα0.9 | CIDRα2.2 | DBLδ1 | CIDRγ8 | DBLε4 | | | | | | | | | | | |
| PFKE07.g1065 | DBLα0.9 | CIDRα2.1 | DBLδ1 | CIDRβ6 | | | | | | | | | | | | |
| PFKE07.g145 | DBLpam1 | DBLpam2 | CIDRpam | DBLpam3 | DBLεpam4 | DBLεpam5 | DBLε10 | | | | | | | | | |
| PFKE07.g1059 | DBLpam1 | DBLpam2 | CIDRpam | DBLpam3 | DBLεpam4 | DBLεpam5 | DBLε10 | DBLε14 | DBLε3 | | | | | | | |
| PFKE07.g74 | CIDRβ1 | | | | | | | | | | | | | | | |
| PFKE07.g414 | DBLδ1 | CIDRβ1 | | | | | | | | | | | | | | |
| PFKE08.g292 | DBLα1.1 | CIDRα1.4 | DBLβ1 | DBLβ3 | DBLδ1 | CIDRβ1 | | | | | | | | | | |
| PFKE08.g355 | DBLα1.1 | CIDRα1.2 | DBLβ11 | DBLγ1 | DBLε1 | DBLγ8 | DBLζ1 | DBLε5 | | | | | | | | |
| PFKE08.g5 | DBLα1.2 | CIDRα1.5 | DBLβ3 | DBLδ1 | CIDRγ2 | | | | | | | | | | | |
| PFKE08.g1 | DBLα1.5 | CIDRδ1 | DBLβ3 | DBLγ2 | DBLβ7 | DBLδ1 | CIDRβ1 | | | | | | | | | |
| PFKE08.g293 | DBLα1.5 | CIDRδ1 | DBLγ13 | DBLγ10 | DBLδ1 | CIDRβ2 | | | | | | | | | | |
| PFKE08.g240 | DBLα1.7 | CIDRα1.4 | DBLβ3 | DBLβ9 | DBLγ11 | DBLβ3 | DBLγ2 | DBLγ4 | DBLα1.6 | CIDRδ2 | DBLβ7 | DBLγ16 | DBLδ5 | CIDRβ3 | DBLβ7 | DBLγ4 |
| PFKE08.g270 | DBLα1.8 | CIDRβ4 | DBLγ2 | DBLδ2 | CIDRγ2 | DBLε13 | DBLζ4 | | | | | | | | | |
| PFKE08.g502 | DBLα1.8 | CIDRβ2 | DBLγ7 | DBLε11 | DBLζ2 | DBLε6 | | | | | | | | | | |
| PFKE08.g239 | DBLα2 | DBLβ12 | DBLζ3 | DBLε6 | DBLδ1 | CIDRγ8 | | | | | | | | | | |
| PFKE08.g501 | DBLα2 | CIDRα1.1 | DBLβ12 | DBLγ4 | DBLδ1 | CIDRβ1 | | | | | | | | | | |
| PFKE08.g4 | DBLα0.1 | CIDRα3.1 | DBLδ1 | CIDRβ1 | | | | | | | | | | | | |
| PFKE08.g8 | DBLα0.1 | CIDRα3.2 | DBLδ1 | CIDRβ5 | | | | | | | | | | | | |
| PFKE08.g11 | DBLα0.1 | CIDRα3.1 | DBLδ1 | CIDRβ1 | | | | | | | | | | | | |
| PFKE08.g214 | DBLα0.1 | CIDRα3.2 | DBLδ1 | CIDRβ4 | | | | | | | | | | | | |
| PFKE08.g290 | DBLα0.1 | CIDRα3.2 | DBLδ1 | CIDRγ7 | | | | | | | | | | | | |
| PFKE08.g291 | DBLα0.1 | CIDRα3.1 | DBLδ1 | CIDRβ1 | | | | | | | | | | | | |
| PFKE08.g424 | DBLα0.1 | CIDRα3.3 | DBLδ1 | CIDRβ1 | | | | | | | | | | | | |
| PFKE08.g3 | DBLα0.11 | CIDRα2.4 | DBLδ1 | CIDRβ1 | | | | | | | | | | | | |
| PFKE08.g9 | DBLα0.11 | CIDRα2.4 | DBLδ1 | CIDRβ1 | | | | | | | | | | | | |
| PFKE08.g250 | DBLα0.11 | CIDRα2.4 | DBLδ1 | CIDRβ5 | | | | | | | | | | | | |
| PFKE08.g357 | DBLα0.11 | CIDRα6 | DBLβ5 | DBLδ5 | CIDRβ4 | DBLγ3 | DBLζ4 | | | | | | | | | |
| PFKE08.g368 | DBLα0.11 | CIDRα2.1 | DBLδ1 | CIDRγ7 | | | | | | | | | | | | |
| PFKE08.g450 | DBLα0.11 | CIDRα2.4 | DBLδ1 | CIDRβ3 | | | | | | | | | | | | |
| PFKE08.g393 | DBLα0.12 | CIDRα2.1 | DBLβ5 | DBLγ11 | DBLζ2 | DBLε3 | | | | | | | | | | |
| PFKE08.g294 | DBLα0.15 | CIDRα3.1 | DBLδ1 | CIDRβ1 | | | | | | | | | | | | |

← New rosetting variant (DC11) - PFKE08varR1

**Rosetting-associated head structure**

DBLα1.5 /6 /8 — CIDRβ /γ /δ

#### PFKE08 (9605)

| | | | | | | | | | |
|---|---|---|---|---|---|---|---|---|---|
| PFKE08.g2 | DBLα0.16 | CIDRα6 | DBLβ5 | DBLδ1 | CIDRγ5 | DBLγ3 | DBLζ3 | DBLε6 | |
| PFKE08.g6 | DBLα0.16 | CIDRα3.4 | DBLδ1 | CIDRγ12 | | | | | |
| PFKE08.g241 | DBLα0.16 | CIDRα3.4 | DBLδ1 | CIDRβ5 | | | | | |
| PFKE08.g229 | DBLα0.17 | CIDRα3.1 | DBLδ1 | CIDRβ1 | | | | | |
| PFKE08.g228 | DBLα0.18 | CIDRα4 | DBLβ5 | DBLγ12 | DBLδ1 | CIDRβ3 | | | |
| PFKE08.g542 | DBLα0.18 | CIDRα6 | DBLβ5 | DBLδ1 | CIDRγ7 | | | | |
| PFKE08.g126 | DBLα0.19 | CIDRα2.5 | DBLδ1 | CIDRβ1 | | | | | |
| PFKE08.g366 | DBLα0.2 | CIDRα3.2 | DBLδ1 | CIDRγ5 | | | | | |
| PFKE08.g238 | DBLα0.21 | CIDRα2.1 | DBLβ2 | DBLγ12 | DBLδ5 | CIDRβ4 | DBLε2 | DBLζ3 | DBLε9 |
| PFKE08.g288 | DBLα0.24 | CIDRα3.1 | DBLδ1 | CIDRγ12 | DBLζ6 | DBLε9 | | | |
| PFKE08.g207 | DBLα0.3 | CIDRα5 | DBLβ5 | DBLγ5 | | | | | |
| PFKE08.g360 | DBLα0.3 | CIDRα4 | DBLβ5 | DBLδ5 | CIDRβ4 | | | | |
| PFKE08.g356 | DBLα0.4 | CIDRα4 | DBLβ5 | DBLδ5 | CIDRβ4 | DBLγ13 | DBLζ5 | DBLε4 | |
| PFKE08.g361 | DBLα0.4 | CIDRα5 | DBLδ1 | CIDRβ1 | DBLγ10 | | | | |
| PFKE08.g362 | DBLα0.4 | CIDRα3.1 | DBLδ1 | CIDRγ4 | | | | | |
| PFKE08.g195 | DBLα0.5 | CIDRα2.1 | DBLδ1 | CIDRβ1 | | | | | |
| PFKE08.g246 | DBLα0.5 | CIDRα2.3 | DBLδ1 | CIDRβ1 | | | | | |
| PFKE08.g496 | DBLα0.6 | CIDRα3.2 | DBLβ5 | DBLγ5 | DBLδ1 | CIDRβ5 | | | |
| PFKE08.g363 | DBLα0.7 | CIDRα2.2 | DBLδ1 | CIDRβ1 | | | | | |
| PFKE08.g297 | DBLα0.8 | CIDRα3.2 | DBLγ5 | DBLδ1 | CIDRβ5 | | | | |
| PFKE08.g12 | DBLα0.9 | CIDRα2.2 | DBLδ1 | CIDRβ1 | | | | | |
| PFKE08.g289 | DBLα0.9 | CIDRα2.1 | DBLδ1 | CIDRβ4 | DBLε2 | DBLε7 | DBLε3 | | |
| PFKE08.g295 | DBLα0.9 | CIDRα2.3 | DBLδ1 | CIDRβ1 | | | | | |
| PFKE08.g296 | DBLα0.9 | CIDRα2.7 | DBLδ1 | CIDRβ5 | | | | | |
| PFKE08.g358 | DBLα0.9 | CIDRα2.3 | DBLδ1 | CIDRγ9 | DBLζ6 | DBLε9 | | | |
| PFKE08.g367 | DBLα0.9 | CIDRα2.3 | DBLδ1 | CIDRβ1 | | | | | |
| PFKE08.g386 | DBLα0.9 | CIDRα2.2 | DBLδ1 | CIDRβ1 | | | | | |
| PFKE08.g486 | DBLα0.9 | CIDRα2.7 | DBLγ11 | DBLζ5 | DBLε4 | | | | |
| PFKE08.g252 | DBLεpam4 | DBLεpam5 | DBLε10 | | | | | | |
| PFKE08.g247 | DBLpam2 | CIDRpam | DBLpam3 | DBLεpam4 | DBLεpam5 | DBLε10 | | | |
| PFKE08.g572 | CIDRα2.3 | DBLδ1 | CIDRβ1 | | | | | | |
| PFKE08.g10 | CIDRα2.5 | DBLβ13 | DBLδ1 | CIDRβ6 | | | | | |
| PFKE08.g20 | CIDRα2.5 | DBLβ13 | DBLδ1 | CIDRβ6 | | | | | |
| PFKE08.g370 | DBLδ1 | CIDRβ1 | | | | | | | |

#### PFKE09 (6816)

| | | | | | | | | | |
|---|---|---|---|---|---|---|---|---|---|
| PFKE09.g227 | DBLα1.1 | CIDRα1.2 | DBLβ11 | DBLγ1 | DBLε1 | DBLγ8 | DBLζ2 | DBLε5 | |
| PFKE09.g471 | DBLα1.1 | CIDRα1.7 | DBLβ6 | DBLγ12 | DBLδ5 | CIDRβ3 | DBLβ9 | DBLγ10 | |
| PFKE09.g331 | DBLα1.2 | CIDRα1.5 | DBLβ3 | DBLγ2 | DBLγ10 | DBLδ1 | CIDRβ1 | | |
| PFKE09.g1 | DBLα1.4 | CIDRα1.7 | DBLγ2 | DBLγ2 | DBLγ4 | DBLδ4 | CIDRγ1 | DBLζ6 | DBLε6 |
| PFKE09.g320 | DBLα1.7 | CIDRα1.8 | DBLβ12 | DBLγ5 | DBLγ4 | DBLδ1 | CIDRγ4 | | |
| PFKE09.g324 | DBLα1.7 | CIDRα1.4 | DBLβ1 | DBLβ7 | DBLγ2 | DBLγ2 | DBLδ1 | CIDRβ1 | |
| PFKE09.g267 | DBLα2 | CIDRα1.8 | DBLβ12 | DBLγ4 | DBLζ5 | DBLε4 | | | |
| PFKE09.g325 | DBLα2 | CIDRα1.1 | DBLβ3 | DBLγ13 | DBLζ2 | DBLε4 | | | |
| PFKE09.g223 | DBLα0.1 | CIDRα3.1 | DBLδ1 | CIDRγ10 | | | | | |
| PFKE09.g333 | DBLα0.1 | CIDRα3.2 | DBLβ4 | | | | | | |
| PFKE09.g426 | DBLα0.1 | CIDRα3.1 | DBLδ1 | CIDRβ1 | | | | | |
| PFKE09.g526 | DBLα0.1 | CIDRα3.2 | DBLδ1 | CIDRγ1 | | | | | |
| PFKE09.g532 | DBLα0.1 | CIDRα3.1 | DBLδ1 | CIDRβ1 | | | | | |
| PFKE09.g234 | DBLα0.1 | CIDRα2.2 | DBLδ1 | CIDRβ1 | | | | | |
| PFKE09.g350 | DBLα0.11 | CIDRα2.4 | DBLδ1 | CIDRγ6 | DBLζ6 | DBLε9 | | | |
| PFKE09.g573 | DBLα0.11 | CIDRα2.4 | DBLδ1 | CIDRγ2 | | | | | |
| PFKE09.g323 | DBLα0.15 | CIDRα3.2 | DBLδ1 | CIDRβ1 | | | | | |
| PFKE09.g410 | DBLα0.15 | CIDRα3.2 | DBLδ1 | CIDRγ7 | | | | | |
| PFKE09.g440 | DBLα0.15 | CIDRα3.2 | DBLδ1 | CIDRβ1 | | | | | |
| PFKE09.g572 | DBLα0.15 | CIDRα2.5 | DBLδ1 | CIDRγ7 | | | | | |
| PFKE09.g230 | DBLα0.16 | CIDRα3.4 | DBLβ13 | DBLδ1 | CIDRβ1 | | | | |
| PFKE09.g3 | DBLα0.17 | CIDRα3.1 | DBLδ1 | CIDRβ1 | | | | | |
| PFKE09.g5 | DBLα0.17 | CIDRα3.3 | DBLδ1 | CIDRβ1 | | | | | |
| PFKE09.g270 | DBLα0.17 | CIDRα3.2 | DBLδ1 | CIDRβ5 | | | | | |
| PFKE09.g95 | DBLα0.18 | CIDRα3.1 | DBLδ1 | CIDRγ1 | DBLε2 | | | | |
| PFKE09.g72 | DBLα0.22 | CIDRα3.1 | DBLδ1 | CIDRγ1 | DBLζ6 | DBLε9 | | | |
| PFKE09.g139 | DBLα0.22 | CIDRα3.2 | DBLδ4 | CIDRδ1 | | | | | |
| PFKE09.g321 | DBLα0.22 | CIDRα3.1 | DBLδ1 | CIDRγ4 | | | | | |
| PFKE09.g332 | DBLα0.3 | CIDRα2.4 | DBLδ1 | CIDRβ1 | | | | | |
| PFKE09.g537 | DBLα0.3 | CIDRα3.2 | DBLδ5 | CIDRβ4 | DBLβ9 | DBLγ2 | | | |
| PFKE09.g2 | DBLα0.5 | CIDRα2.1 | DBLδ1 | CIDRβ1 | | | | | |
| PFKE09.g142 | DBLα0.5 | CIDRα2.5 | | | | | | | |
| PFKE09.g269 | DBLα0.5 | CIDRα2.6 | DBLδ1 | CIDRβ5 | | | | | |
| PFKE09.g322 | DBLα0.5 | CIDRα2.3 | DBLδ1 | CIDRβ1 | | | | | |
| PFKE09.g340 | DBLα0.5 | CIDRα2.9 | DBLδ1 | CIDRβ3 | | | | | |
| PFKE09.g73 | DBLα0.6 | CIDRα4 | DBLδ1 | CIDRβ4 | DBLε13 | | | | |
| PFKE09.g399 | DBLα0.6 | CIDRα3.1 | DBLδ1 | CIDRβ1 | | | | | |

| | | | | | | | | | |
|---|---|---|---|---|---|---|---|---|---|
| PFKE09.g268 | DBLα0.7 | CIDRα3.5 | DBLδ1 | CIDRβ6 | | | | | |
| PFKE09.g75 | DBLα0.8 | CIDRα5 | DBLβ5 | DBLδ1 | CIDRγ9 | DBLε2 | DBLζ3 | DBLε3 | |
| PFKE09.g357 | DBLα0.9 | CIDRα2.2 | DBLδ1 | CIDRγ7 | | | | | |
| PFKE09.g372 | DBLα0.9 | CIDRα2.4 | DBLδ1 | CIDRβ1 | | | | | |
| PFKE09.g575 | DBLα0.9 | CIDRα3.2 | DBLδ1 | CIDRβ6 | | | | | |
| PFKE09.g369 | DBLpam1 | CIDRpam | | | | | | | |
| PFKE09.g159 | CIDRα1.8 | DBLβ12 | DBLγ6 | | | | | | |
| PFKE09.g368 | CIDRα1.8 | DBLβ12 | DBLγ6 | | | | | | |
| PFKE09.g56 | CIDRγ6 | DBLζ6 | DBLε9 | | | | | | |
| PFKE09.g110 | DBLδ1 | | | | | | | | |
| PFKE09.g500 | DBLδ1 | CIDRβ1 | | | | | | | |
| PFKE09.g563 | DBLδ7 | CIDRβ1 | | | | | | | |
| PFKE10.g6 | DBLα1.1 | CIDRα1.4 | DBLγ9 | DBLβ3 | DBLδ1 | CIDRβ1 | | | |
| PFKE10.g237 | DBLα1.1 | CIDRα1.2 | DBLβ11 | DBLγ1 | DBLε1 | DBLγ8 | DBLζ1 | DBLε5 | |
| PFKE10.g399 | DBLα1.1 | CIDRα1.7 | DBLβ3 | DBLγ2 | DBLγ4 | DBLγ2 | DBLδ1 | CIDRγ1 | |
| PFKE10.g1 | DBLα1.2 | CIDRα1.5 | DBLβ6 | DBLγ16 | DBLδ5 | CIDRβ3 | DBLβ7 | DBLγ2 | |
| PFKE10.g363 | DBLα1.2 | CIDRα1.5 | DBLβ3 | DBLγ2 | DBLγ4 | | | | |
| PFKE10.g462 | DBLα1.2 | CIDRα1.7 | DBLβ1 | DBLγ2 | DBLγ4 | DBLγ2 | DBLζ3 | DBLε6 | |
| PFKE10.g435 | DBLα1.3 | DBLε8 | | | | | | | |
| PFKE10.g472 | DBLα1.6 | CIDRγ3 | DBLγ6 | DBLδ2 | CIDRγ6 | DBLε13 | DBLζ5 | DBLε4 | |
| PFKE10.varR1 | DBLα1.8 | CIDRγ3 | DBLγ7 | DBLε11 | DBLζ3 | DBLε8 | | | |
| PFKE10.g243 | DBLα1.8 | CIDRγ3 | DBLγ13 | DBLγ10 | DBLδ1 | CIDRβ5 | | | |
| PFKE10.g239 | DBLα2 | CIDRα1.8 | DBLβ12 | DBLζ3 | DBLε5 | DBLδ1 | CIDRβ1 | | |
| PFKE10.g283 | DBLα2 | CIDRα1.1 | DBLβ4 | DBLγ4 | DBLδ1 | CIDRβ6 | | | |
| PFKE10.g248 | CIDRα1.7 | DBLα0.12 | DBLγ2 | | | | | | |
| PFKE10.g27 | DBLα0.1 | CIDRα3.5 | | | | | | | |
| PFKE10.g217 | DBLα0.1 | CIDRα3.1 | DBLδ6 | | | | | | |
| PFKE10.g223 | DBLα0.1 | CIDRα3.2 | DBLδ1 | CIDRγ1 | | | | | |
| PFKE10.g286 | DBLα0.1 | CIDRα3.1 | DBLδ1 | CIDRγ4 | | | | | |
| PFKE10.g320 | DBLα0.1 | CIDRα3.1 | DBLδ1 | CIDRβ1 | | | | | |
| PFKE10.g14 | DBLα0.1 | CIDRα2.2 | DBLδ1 | CIDRβ6 | | | | | |
| PFKE10.g247 | DBLα0.1 | CIDRα2.2 | DBLδ1 | CIDRγ1 | DBLε2 | DBLζ3 | DBLε3 | | |
| PFKE10.g3 | DBLα0.11 | CIDRα2.4 | DBLβ5 | DBLδ1 | CIDRβ1 | | | | |
| PFKE10.g8 | DBLα0.11 | CIDRα2.1 | DBLδ1 | CIDRβ1 | | | | | |
| PFKE10.g48 | DBLα0.11 | CIDRα2.4 | DBLδ1 | CIDRβ1 | | | | | |
| PFKE10.g246 | DBLα0.11 | CIDRα2.4 | DBLδ1 | CIDRβ5 | | | | | |
| PFKE10.g288 | DBLα0.11 | CIDRα2.4 | DBLδ1 | CIDRγ4 | | | | | |
| PFKE10.g525 | DBLα0.11 | CIDRα2.4 | DBLδ1 | CIDRβ1 | | | | | |
| PFKE10.g473 | DBLα0.13 | CIDRα2.3 | DBLδ1 | | | | | | |
| PFKE10.g10 | DBLα0.15 | CIDRα3.1 | DBLδ1 | CIDRβ5 | | | | | |
| PFKE10.g103 | DBLα0.15 | CIDRα3.2 | DBLδ1 | CIDRβ1 | | | | | |
| PFKE10.g130 | DBLα0.15 | CIDRα3.2 | DBLδ1 | CIDRβ7 | | | | | |
| PFKE10.g280 | DBLα0.15 | CIDRα3.2 | DBLβ5 | DBLδ8 | CIDRβ2 | DBLγ3 | DBLζ4 | | |
| PFKE10.g484 | DBLα0.15 | CIDRα3.2 | DBLδ1 | CIDRβ1 | | | | | |
| PFKE10.g13 | DBLα0.17 | CIDRα3.1 | | | | | | | |
| PFKE10.g72 | DBLα0.17 | CIDRα3.1 | DBLδ1 | CIDRβ5 | | | | | |
| PFKE10.g289 | DBLα0.17 | CIDRα3.3 | DBLδ1 | CIDRβ6 | | | | | |
| PFKE10.g249 | DBLα0.18 | CIDRα4 | DBLβ5 | DBLγ18 | DBLε8 | | | | |
| PFKE10.g284 | DBLα0.18 | CIDRα5 | DBLβ5 | DBLδ1 | CIDRβ1 | | | | |
| PFKE10.g9 | DBLα0.2 | CIDRα3.2 | DBLδ1 | CIDRβ1 | | | | | |
| PFKE10.g287 | DBLα0.22 | CIDRα3.1 | DBLδ1 | CIDRβ1 | | | | | |
| PFKE10.g291 | DBLα0.23 | CIDRα6 | DBLβ5 | DBLγ2 | DBLδ3 | CIDRγ2 | DBLε2 | DBLε7 | DBLε3 |
| PFKE10.g267 | DBLα0.3 | CIDRα4 | DBLβ5 | DBLδ5 | CIDRβ4 | | | | |
| PFKE10.g282 | DBLα0.4 | CIDRα5 | DBLβ6 | DBLγ9 | | | | | |
| PFKE10.g7 | DBLα0.5 | CIDRα2.3 | DBLδ1 | CIDRγ2 | DBLε2 | DBLε7 | DBLε3 | | |
| PFKE10.g11 | DBLα0.5 | CIDRα2.3 | DBLδ1 | CIDRβ1 | DBLα0.12 | CIDRα2.11 | DBLε2 | DBLζ3 | DBLε6 |
| PFKE10.g15 | DBLα0.5 | CIDRα2.6 | DBLδ1 | CIDRβ1 | | | | | |
| PFKE10.g299 | DBLα0.5 | CIDRα2.3 | DBLδ1 | CIDRγ1 | DBLγ3 | DBLζ4 | | | |
| PFKE10.g365 | DBLα0.5 | CIDRα2.1 | DBLβ5 | DBLδ1 | CIDRβ5 | | | | |
| PFKE10.g4 | DBLα0.6 | CIDRα3.1 | DBLβ5 | DBLγ5 | DBLγ17 | DBLδ1 | CIDRβ6 | | |
| PFKE10.g20 | DBLα0.6 | CIDRα3.1 | | | | | | | |
| PFKE10.g181 | DBLα0.6 | CIDRα5 | | | | | | | |
| PFKE10.g524 | DBLα0.6 | CIDRα3.3 | DBLβ5 | DBLδ1 | CIDRβ1 | | | | |
| PFKE10.g16 | DBLα0.8 | CIDRα5 | DBLδ1 | CIDRβ1 | | | | | |
| PFKE10.g21 | DBLα0.9 | CIDRα2.1 | DBLδ1 | CIDRβ1 | | | | | |
| PFKE10.g285 | DBLα0.9 | CIDRα2.2 | DBLδ1 | CIDRβ1 | | | | | |
| PFKE10.g389 | DBLpam1 | DBLpam2 | CIDRpam | | | | | | |
| PFKE10.g517 | DBLpam2 | CIDRpam | DBLpam3 | DBLε4 | DBLεpam5 | DBLε3 | DBLε10 | | |
| PFKE10.g2 | CIDRα1.7 | DBLγ2 | DBLγ2 | | | | | | |
| PFKE10.g252 | CIDRα2.4 | DBLδ1 | CIDRβ1 | | | | | | |
| PFKE10.g364 | CIDRα2.4 | DBLδ1 | CIDRβ6 | | | | | | |

New rosetting variant (DC11) - PFKE10varR1

**Rosetting-associated head structure**

DBLα1.5 /6 /8 — CIDRβ /γ /δ

### PFKE10 (11019)

| | | | | | | | | |
|---|---|---|---|---|---|---|---|---|
| PFKE10.g81 | CIDRα3.1 | DBLδ1 | CIDRβ1 | | | | | |
| PFKE10.g416 | CIDRα5 | DBLδ1 | CIDRβ6 | | | | | |
| PFKE10.g292 | DBLβ5 | DBLβ3 | DBLδ1 | CIDRβ1 | | | | |
| PFKE10.g245 | DBLβ8 | DBLβ5 | DBLδ1 | CIDRβ1 | | | | |
| PFKE10.g17 | DBLδ1 | CIDRβ1 | | | | | | |
| PFKE10.g71 | DBLδ1 | CIDRβ1 | | | | | | |
| PFKE10.g264 | DBLδ1 | CIDRβ1 | | | | | | |
| PFKE10.g431 | DBLε9 | | | | | | | |
| PFKE11.g431 | DBLα1.1 | CIDRα1.7 | DBLβ3 | DBLγ6 | DBLγ9 | DBLδ1 | CIDRγ5 | |
| PFKE11.g554 | DBLα1.1 | CIDRα1.2 | DBLβ11 | DBLγ1 | DBLε1 | | | |
| PFKE11.g448 | DBLα1.2 | CIDRα1.5 | DBLβ6 | DBLγ2 | DBLγ4 | DBLδ1 | CIDRβ6 | |
| PFKE11.g479 | DBLα1.2 | CIDRα1.5 | DBLγ11 | | | | | |
| PFKE11.g604 | DBLα1.2 | CIDRα1.5 | DBLβ3 | DBLγ10 | DBLδ1 | CIDRβ5 | | |
| PFKE11.g54 | DBLα1.3 | DBLε8 | | | | | | |
| PFKE11.g294 | DBLα1.4 | CIDRα1.4 | DBLβ1 | DBLβ7 | DBLγ14 | DBLδ1 | CIDRβ6 | |
| PFKE11.g553 | DBLα1.4 | CIDRα1.3 | DBLβ1 | DBLγ15 | DBLε1 | | | |
| PFKE11.g500 | DBLα1.5 | CIDRδ1 | DBLγ11 | DBLγ10 | DBLδ1 | CIDRβ1 | | |
| PFKE11.g6 | DBLα1.6 | CIDRδ2 | DBLβ6 | DBLγ14 | DBLζ5 | DBLε4 | | |
| PFKE11.g264 | DBLα1.6 | CIDRγ3 | DBLγ7 | DBLδ5 | CIDRβ3 | DBLβ6 | DBLε13 | |
| PFKE11.g2 | DBLα1.7 | CIDRα1.4 | DBLβ1 | DBLγ6 | DBLγ10 | DBLδ5 | CIDRβ4 | |
| PFKE11.g536 | DBLα1.7 | CIDRα1.4 | DBLγ2 | DBLβ6 | DBLδ1 | CIDRβ1 | | |
| PFKE11.g7 | DBLα2 | CIDRα1.7 | DBLγ2 | DBLγ2 | DBLγ4 | DBLζ3 | DBLε6 | |
| PFKE11.g224 | DBLα2 | CIDRα1.1 | DBLβ12 | DBLγ6 | | | | |
| PFKE11.g359 | DBLα2 | CIDRα1.1 | DBLβ12 | DBLγ6 | DBLδ1 | CIDRγ12 | | |
| PFKE11.g375 | DBLα2 | CIDRα1.1 | DBLβ12 | DBLγ11 | DBLζ3 | DBLε12 | | |
| PFKE11.g456 | DBLα2 | CIDRα1.1 | DBLβ12 | DBLγ6 | DBLγ17 | DBLζ5 | DBLε4 | |
| PFKE11.g486 | DBLα2 | CIDRα1.1 | DBLβ12 | DBLγ6 | | | | |
| PFKE11.g11 | DBLα0.1 | CIDRα3.1 | DBLδ1 | CIDRβ1 | | | | |
| PFKE11.g137 | DBLα0.1 | CIDRα3.1 | DBLδ1 | CIDRβ1 | | | | |
| PFKE11.g159 | DBLα0.1 | CIDRα3.1 | DBLδ1 | CIDRγ6 | | | | |
| PFKE11.g217 | DBLα0.1 | CIDRα3.1 | DBLδ1 | CIDRβ3 | | | | |
| PFKE11.g270 | DBLα0.1 | CIDRα3.2 | DBLδ1 | CIDRβ5 | | | | |
| PFKE11.g306 | DBLα0.1 | CIDRα3.2 | DBLδ1 | CIDRβ6 | | | | |
| PFKE11.g338 | DBLα0.1 | CIDRα3.1 | DBLδ1 | DBLα0.11 | CIDRα2.2 | DBLδ1 | CIDRβ3 | |
| PFKE11.g360 | DBLα0.1 | CIDRα3.1 | DBLδ1 | CIDRβ1 | | | | |
| PFKE11.g466 | DBLα0.1 | CIDRα3.2 | DBLδ1 | CIDRβ5 | | | | |
| PFKE11.g671 | DBLα0.1 | CIDRα2.4 | DBLδ1 | CIDRβ5 | | | | |
| PFKE11.g619 | DBLα0.1 | CIDRα2.2 | DBLβ5 | DBLδ1 | CIDRβ1 | | | |
| PFKE11.g293 | DBLα0.11 | CIDRα2.4 | DBLγ4 | DBLδ4 | CIDRγ1 | DBLε2 | DBLε7 | DBLε3 |
| PFKE11.g350 | DBLα0.11 | CIDRα2.4 | DBLδ1 | CIDRγ11 | | | | |
| PFKE11.g450 | DBLα0.11 | CIDRα2.2 | DBLδ1 | CIDRβ1 | | | | |
| PFKE11.g451 | DBLα0.11 | CIDRα2.1 | DBLδ1 | CIDRβ1 | | | | |
| PFKE11.g362 | DBLα0.12 | CIDRα2.2 | DBLδ1 | CIDRβ1 | | | | |
| PFKE11.g366 | DBLα0.12 | CIDRα2.11 | DBLδ1 | CIDRβ1 | | | | |
| PFKE11.g436 | DBLα0.12 | CIDRα2.2 | DBLδ1 | CIDRβ1 | | | | |
| PFKE11.g698 | DBLα0.12 | CIDRα2.1 | DBLγ2 | DBLδ1 | CIDRβ1 | | | |
| PFKE11.g9 | DBLα0.13 | CIDRα2.1 | DBLγ2 | DBLδ1 | CIDRβ1 | | | |
| PFKE11.g10 | DBLα0.13 | CIDRα2.3 | DBLβ5 | DBLδ1 | CIDRβ1 | | | |
| PFKE11.g295 | DBLα0.13 | CIDRα2.5 | DBLβ5 | DBLδ1 | CIDRβ5 | | | |
| PFKE11.g355 | DBLα0.13 | CIDRα2.3 | DBLδ1 | CIDRβ6 | | | | |
| PFKE11.g297 | DBLα0.15 | CIDRα3.2 | DBLδ1 | CIDRβ1 | | | | |
| PFKE11.g353 | DBLα0.15 | CIDRα3.2 | DBLδ1 | CIDRγ1 | DBLζ6 | DBLε9 | | |
| PFKE11.g361 | DBLα0.15 | CIDRα3.1 | DBLδ1 | CIDRβ1 | | | | |
| PFKE11.g19 | DBLα0.16 | CIDRα3.4 | DBLδ1 | CIDRβ6 | | | | |
| PFKE11.g33 | DBLα0.16 | CIDRα3.4 | DBLδ1 | CIDRγ4 | | | | |
| PFKE11.g480 | DBLα0.16 | CIDRα3.4 | DBLδ1 | CIDRβ1 | | | | |
| PFKE11.g58 | DBLα0.17 | CIDRα3.1 | DBLδ1 | CIDRβ1 | | | | |
| PFKE11.g298 | DBLα0.17 | CIDRα3.1 | DBLδ1 | CIDRβ1 | | | | |
| PFKE11.g351 | DBLα0.17 | CIDRα3.3 | DBLδ1 | CIDRβ5 | | | | |
| PFKE11.g367 | DBLα0.17 | CIDRα3.1 | DBLδ1 | CIDRγ11 | | | | |
| PFKE11.g20 | DBLα0.18 | CIDRα6 | DBLβ4 | | | | | |
| PFKE11.g570 | DBLα0.18 | CIDRα6 | DBLβ5 | DBLδ4 | CIDRδ1 | DBLβ6 | | |
| PFKE11.g8 | DBLα0.19 | CIDRα2.4 | DBLδ1 | CIDRγ11 | | | | |
| PFKE11.g5 | DBLα0.2 | CIDRα3.1 | DBLδ1 | CIDRβ5 | | | | |
| PFKE11.g15 | DBLα0.2 | CIDRα3.1 | DBLδ1 | | | | | |
| PFKE11.g354 | DBLα0.22 | CIDRα3.3 | DBLδ1 | CIDRβ1 | | | | |
| PFKE11.g697 | DBLα0.23 | CIDRα6 | DBLδ1 | CIDRβ1 | | | | |
| PFKE11.g214 | DBLα0.3 | CIDRα5 | DBLβ4 | DBLγ2 | DBLδ9 | | | |
| PFKE11.g444 | DBLα0.3 | CIDRα3.1 | DBLγ5 | DBLδ1 | CIDRβ6 | | | |
| PFKE11.g518 | DBLα0.3 | CIDRα2.1 | DBLγ11 | DBLζ2 | DBLε4 | | | |
| PFKE11.g649 | DBLα0.3 | CIDRα3.2 | DBLδ1 | CIDRγ5 | | | | |

New rosetting variant (DC15) - PFKE11varR1

**Rosetting-associated head structure**

DBLα1.5 /6 /8 | CIDRβ /γ /δ

### PFKE11 (9775)

| | | | | | | | | | | |
|---|---|---|---|---|---|---|---|---|---|---|
| PFKE11.g332 | DBLα0.4 | CIDRα3.1 | DBLδ1 | CIDRβ1 | | | | | | |
| PFKE11.g345 | DBLα0.4 | CIDRα6 | DBLβ5 | DBLδ5 | CIDRβ4 | | | | | |
| PFKE11.g296 | DBLα0.5 | CIDRα2.5 | DBLβ13 | DBLδ1 | CIDRβ5 | | | | | |
| PFKE11.g352 | DBLα0.5 | CIDRα2.5 | DBLδ1 | CIDRβ6 | | | | | | |
| PFKE11.g709 | DBLα0.5 | CIDRα2.5 | DBLβ5 | DBLδ1 | CIDRβ1 | | | | | |
| PFKE11.g18 | DBLα0.7 | CIDRα2.2 | DBLδ1 | CIDRγ4 | | | | | | |
| PFKE11.g445 | DBLα0.7 | CIDRα4 | DBLδ1 | CIDRβ1 | | | | | | |
| PFKE11.g517 | DBLα0.8 | CIDRα5 | DBLβ5 | DBLγ10 | | | | | | |
| PFKE11.g1 | DBLα0.9 | CIDRα2.1 | DBLβ2 | DBLδ1 | CIDRγ4 | DBLγ16 | DBLζ5 | DBLε11 | DBLε3 | |
| PFKE11.g73 | DBLα0.9 | CIDRα2.4 | DBLδ4 | CIDRδ1 | | | | | | |
| PFKE11.g140 | DBLα0.9 | CIDRα2.4 | DBLγ9 | DBLδ1 | CIDRγ7 | | | | | |
| PFKE11.g312 | DBLα0.9 | CIDRα2.11 | DBLδ1 | CIDRβ1 | | | | | | |
| PFKE11.g484 | DBLα0.9 | CIDRα2.7 | DBLδ1 | CIDRβ3 | | | | | | |
| PFKE11.g647 | DBLα0.9 | CIDRα6 | DBLβ6 | DBLγ17 | DBLδ1 | CIDRβ1 | | | | |
| PFKE11.g708 | DBLα0.9 | CIDRα2.2 | DBLε2 | DBLζ3 | DBLε12 | | | | | |
| PFKE11.g302 | CIDRα2.5 | DBLδ1 | CIDRβ1 | | | | | | | |
| PFKE11.g459 | CIDRα2.7 | DBLδ1 | CIDRβ1 | | | | | | | |
| PFKE11.g12 | DBLβ4 | DBLβ10 | DBLδ1 | CIDRβ1 | | | | | | |
| PFKE11.g447 | DBLβ6 | DBLβ8 | DBLδ1 | CIDRγ4 | | | | | | |
| PFKE11.g76 | DBLδ1 | CIDRβ5 | | | | | | | | |
| PFKE11.g91 | DBLδ1 | CIDRγ4 | | | | | | | | |
| PFKE11.g538 | DBLδ1 | CIDRβ1 | | | | | | | | |
| PFKE11.g349 | DBLγ2 | DBLγ4 | DBLγ2 | DBLγ12 | DBLδ1 | CIDRβ1 | | | | |
| PFKE11.g552 | DBLζ2 | DBLε5 | | | | | | | | |
| PFKE12.g232 | DBLα1.1 | CIDRα1.2 | DBLβ11 | DBLγ1 | DBLε1 | DBLγ8 | DBLζ2 | DBLε5 | | |
| PFKE12.g307 | DBLα1.1 | CIDRα1.4 | DBLβ12 | DBLγ6 | DBLγ10 | | | | | |
| PFKE12.g5 | DBLα1.2 | CIDRα1.4 | DBLβ12 | DBLγ5 | DBLγ2 | DBLδ1 | CIDRβ1 | | | |
| PFKE12.g251 | DBLα1.2 | CIDRα1.5 | DBLβ6 | DBLδ4 | CIDRδ1 | DBLβ9 | | | | |
| PFKE12.g313 | DBLα1.2 | CIDRα1.4 | DBLβ7 | DBLβ7 | DBLγ2 | DBLγ4 | DBLδ1 | CIDRγ2 | | |
| PFKE12.g23 | DBLα1.3 | DBLε8 | | | | | | | | |
| PFKE12.g2 | DBLα1.4 | CIDRα1.5 | DBLβ3 | DBLβ6 | DBLδ1 | CIDRβ1 | | | | |
| PFKE12.g386 | DBLα1.4 | CIDRα1.7 | DBLβ1 | DBLγ11 | DBLγ2 | DBLγ4 | DBLδ1 | | | |
| PFKE12.g11 | DBLα1.6 | CIDRγ3 | DBLγ12 | DBLδ5 | CIDRβ4 | DBLβ7 | | | | |
| PFKE12.g233 | DBLα1.6 | CIDRδ1 | DBLβ3 | DBLγ2 | DBLγ4 | | | | | |
| PFKE12.g497 | DBLα1.6 | CIDRδ1 | DBLγ12 | DBLδ4 | CIDRγ2 | DBLβ6 | | | | |
| PFKE12.g14 | DBLα1.7 | CIDRα1.4 | DBLγ2 | DBLβ7 | DBLγ2 | | | | | |
| PFKE12.g312 | DBLβ5 | DBLβ3 | DBLδ1 | CIDRβ1 | DBLα2 | CIDRα1.8 | DBLβ12 | DBLγ4 | DBLζ5 | DBLε4 |
| PFKE12.g10 | DBLα0.1 | CIDRα3.3 | DBLδ1 | CIDRβ1 | DBLγ10 | | | | | |
| PFKE12.g18 | DBLα0.1 | CIDRα3.2 | DBLδ1 | CIDRβ1 | | | | | | |
| PFKE12.g89 | DBLα0.1 | CIDRα3.1 | DBLδ1 | CIDRγ2 | | | | | | |
| PFKE12.g150 | DBLα0.1 | CIDRα3.1 | DBLδ1 | CIDRβ5 | | | | | | |
| PFKE12.g234 | DBLα0.1 | CIDRα3.1 | DBLδ1 | CIDRβ1 | | | | | | |
| PFKE12.g272 | DBLα0.1 | CIDRα4 | DBLδ1 | CIDRβ5 | | | | | | |
| PFKE12.g305 | DBLα0.1 | CIDRα3.2 | DBLδ1 | CIDRγ4 | | | | | | |
| PFKE12.g310 | DBLα0.1 | CIDRα3.1 | DBLδ1 | CIDRβ1 | | | | | | |
| PFKE12.g311 | DBLα0.1 | CIDRα3.1 | DBLδ1 | CIDRβ1 | | | | | | |
| PFKE12.g465 | DBLα0.1 | CIDRα3.2 | DBLδ1 | CIDRβ1 | | | | | | |
| PFKE12.g274 | DBLα0.1 | CIDRα2.2 | DBLδ1 | CIDRβ1 | | | | | | |
| PFKE12.g25 | DBLα0.11 | CIDRα2.1 | DBLδ1 | CIDRγ5 | | | | | | |
| PFKE12.g73 | DBLα0.11 | CIDRα2.1 | DBLδ1 | CIDRβ1 | | | | | | |
| PFKE12.g269 | DBLα0.11 | CIDRα2.4 | DBLβ5 | DBLδ1 | CIDRβ6 | | | | | |
| PFKE12.g356 | DBLα0.11 | CIDRα2.4 | DBLδ1 | CIDRβ1 | | | | | | |
| PFKE12.g383 | DBLα0.11 | CIDRα2.4 | DBLδ1 | CIDRβ6 | | | | | | |
| PFKE12.g1 | DBLα0.12 | CIDRα2.1 | DBLβ5 | DBLδ1 | CIDRβ1 | | | | | |
| PFKE12.g8 | DBLα0.12 | CIDRα2.1 | DBLδ1 | CIDRβ1 | | | | | | |
| PFKE12.g347 | DBLα0.13 | CIDRα2.9 | DBLδ1 | CIDRβ1 | | | | | | |
| PFKE12.g106 | DBLα0.14 | CIDRα4 | DBLδ1 | CIDRγ1 | | | | | | |
| PFKE12.g170 | DBLα0.17 | CIDRα3.2 | DBLδ1 | CIDRβ1 | | | | | | |
| PFKE12.g271 | DBLα0.17 | CIDRα3.2 | DBLδ1 | CIDRβ1 | | | | | | |
| PFKE12.g498 | DBLα0.17 | CIDRα3.1 | DBLδ1 | CIDRβ1 | | | | | | |
| PFKE12.g38 | DBLα0.18 | CIDRα4 | DBLβ5 | DBLδ1 | CIDRβ6 | | | | | |
| PFKE12.g105 | DBLα0.18 | CIDRα4 | DBLβ5 | | | | | | | |
| PFKE12.g268 | DBLα0.18 | CIDRα6 | DBLβ4 | DBLγ12 | DBLδ5 | CIDRβ3 | DBLζ1 | DBLε14 | | |
| PFKE12.g338 | DBLα0.18 | CIDRα4 | DBLβ5 | DBLδ1 | CIDRβ6 | | | | | |
| PFKE12.g28 | DBLα0.19 | CIDRα2.6 | DBLδ1 | CIDRβ1 | | | | | | |
| PFKE12.g7 | DBLα0.21 | CIDRα2.1 | DBLβ2 | DBLγ10 | DBLδ9 | CIDRγ2 | | | | |
| PFKE12.g210 | DBLα0.4 | CIDRα5 | DBLβ5 | DBLγ5 | DBLδ1 | CIDRβ1 | | | | |
| PFKE12.g324 | DBLα0.4 | CIDRα6 | DBLδ1 | CIDRβ1 | | | | | | |
| PFKE12.g303 | DBLα0.5 | CIDRα2.3 | DBLδ1 | CIDRγ4 | DBLε2 | DBLε7 | DBLε3 | | | |
| PFKE12.g375 | DBLα0.6 | CIDRα5 | DBLβ5 | DBLγ11 | | | | | | |
| PFKE12.g48 | DBLα0.8 | CIDRα3.4 | DBLδ1 | CIDRγ4 | | | | | | |

**PFKE12 (9215)**

Rosetting-associated head structure

DBLα1.5 /6 /8 — CIDRβ /γ /δ

| | | | | | | | | | |
|---|---|---|---|---|---|---|---|---|---|
| PFKE12.g74 | DBLα0.8 | CIDRα3.4 | DBLδ1 | CIDRβ1 | | | | | |
| PFKE12.g250 | DBLα0.8 | CIDRα4 | DBLδ1 | CIDRβ1 | | | | | |
| PFKE12.g263 | DBLα0.8 | CIDRα3.4 | DBLδ1 | CIDRβ1 | | | | | |
| PFKE12.g85 | DBLα0.9 | CIDRα2.1 | DBLδ1 | CIDRβ5 | | | | | |
| PFKE12.g306 | DBLα0.9 | CIDRα2.2 | DBLδ1 | CIDRγ11 | DBLγ3 | DBLζ4 | | | |
| PFKE12.g316 | DBLα0.9 | CIDRα2.7 | DBLδ1 | CIDRβ1 | | | | | |
| PFKE12.g77 | DBLpam1 | DBLpam2 | CIDRpam | DBLpam3 | DBLεpam4 | DBLεpam5 | DBLε10 | DBLε14 | DBLε3 |
| PFKE12.g429 | DBLpam1 | DBLpam2 | CIDRpam | DBLpam3 | DBLεpam4 | DBLεpam5 | DBLε10 | | |
| PFKE12.g13 | CIDRα2.1 | DBLδ1 | CIDRγ9 | DBLζ6 | DBLε6 | | | | |
| PFKE12.g321 | CIDRα2.1 | DBLδ1 | CIDRβ1 | | | | | | |
| PFKE12.g157 | CIDRα4 | DBLδ1 | CIDRβ1 | | | | | | |
| PFKE12.g3 | DBLβ13 | DBLβ5 | DBLβ5 | DBLδ1 | CIDRβ1 | | | | |
| PFKE12.g12 | DBLβ8 | CIDRα4 | DBLδ1 | CIDRγ1 | DBLζ6 | DBLε9 | | | |
| PFKE12.g291 | DBLδ1 | CIDRγ4 | | | | | | | |
| PFKE12.g314 | DBLδ1 | CIDRβ1 | | | | | | | |
| PC0053-C.g342 | DBLα1.1 | CIDRα1.2 | DBLβ11 | DBLγ1 | DBLε1 | DBLγ8 | DBLζ1 | DBLε5 | |
| PC0053-C.g157 | DBLα1.2 | CIDRα1.5 | DBLβ9 | DBLγ11 | DBLδ1 | CIDRγ8 | | | |
| PC0053-C.g687 | DBLα1.2 | CIDRα1.5 | DBLγ11 | DBLδ4 | CIDRγ9 | DBLε13 | DBLζ5 | DBLε4 | |
| PC0053-C.g20 | DBLα1.4 | CIDRγ3 | DBLβ7 | DBLγ12 | DBLδ3 | CIDRγ2 | DBLζ6 | DBLε9 | |
| PC0053-C.g325 | DBLα1.5 | CIDRδ1 | DBLγ12 | DBLδ5 | CIDRβ3 | DBLβ9 | | | |
| PC0053-C.g188 | DBLα1.6 | CIDRγ3 | DBLγ7 | DBLε11 | DBLα1.3 | DBLε8 | | | |
| PC0053-C.g741 | DBLα1.6 | CIDRγ3 | DBLγ15 | DBLε1 | DBLε11 | DBLζ2 | DBLε6 | | |
| PC0053-C.g111 | DBLα1.7 | CIDRα1.4 | DBLγ2 | DBLγ4 | DBLγ11 | DBLδ1 | CIDRβ1 | | |
| PC0053-C.g96 | DBLα2 | CIDRα1.1 | DBLβ12 | DBLγ6 | DBLγ14 | DBLζ5 | DBLε4 | | |
| PC0053-C.g586 | DBLα2 | CIDRα1.4 | DBLβ3 | DBLγ14 | DBLζ3 | DBLε12 | | | |
| PC0053-C.g211 | DBLα0.1 | CIDRα3.3 | DBLδ1 | CIDRγ1 | DBLε2 | DBLε7 | DBLε3 | | |
| PC0053-C.g243 | DBLα0.1 | CIDRα3.1 | DBLδ1 | CIDRβ2 | DBLε13 | | | | |
| PC0053-C.g567 | DBLα0.1 | CIDRα3.1 | DBLβ5 | DBLγ5 | | | | | |
| PC0053-C.g576 | DBLα0.1 | CIDRα3.1 | DBLδ1 | | | | | | |
| PC0053-C.g594 | DBLα0.1 | CIDRα3.2 | DBLδ1 | CIDRβ1 | | | | | |
| PC0053-C.g599 | DBLα0.1 | CIDRα3.1 | DBLδ1 | CIDRβ5 | | | | | |
| PC0053-C.g606 | DBLα0.1 | CIDRα3.2 | DBLδ1 | CIDRβ5 | | | | | |
| PC0053-C.g619 | DBLα0.1 | CIDRα3.1 | DBLδ1 | CIDRγ12 | | | | | |
| PC0053-C.g648 | DBLα0.1 | CIDRα3.1 | DBLδ1 | CIDRβ1 | | | | | |
| PC0053-C.g284 | DBLα0.11 | CIDRα2.4 | DBLδ3 | CIDRγ2 | DBLε2 | DBLε7 | DBLε3 | | |
| PC0053-C.g685 | DBLα0.11 | CIDRα2.4 | DBLδ1 | CIDRβ1 | | | | | |
| PC0053-C.g486 | DBLα0.12 | CIDRα2.1 | DBLγ11 | DBLζ3 | DBLε12 | | | | |
| PC0053-C.g675 | DBLα0.12 | CIDRα2.2 | DBLδ1 | CIDRβ1 | | | | | |
| PC0053-C.g624 | DBLα0.13 | CIDRα2.3 | DBLδ1 | CIDRβ1 | | | | | |
| PC0053-C.g684 | DBLα0.13 | CIDRα2.6 | DBLδ1 | CIDRβ1 | | | | | |
| PC0053-C.g10 | DBLα0.15 | CIDRα3.2 | | | | | | | |
| PC0053-C.g472 | DBLα0.16 | CIDRα3.4 | DBLγ2 | DBLδ1 | CIDRβ1 | | | | |
| PC0053-C.g587 | DBLα0.16 | CIDRα3.4 | DBLδ1 | CIDRγ4 | | | | | |
| PC0053-C.g649 | DBLα0.16 | CIDRα3.4 | DBLδ1 | CIDRβ1 | | | | | |
| PC0053-C.g686 | DBLα0.16 | CIDRα6 | DBLδ1 | CIDRβ5 | | | | | |
| PC0053-C.g688 | DBLα0.16 | CIDRα3.4 | DBLδ1 | CIDRγ7 | | | | | |
| PC0053-C.g475 | DBLα0.17 | CIDRα3.1 | DBLδ1 | CIDRγ6 | | | | | |
| PC0053-C.g664 | DBLα0.17 | CIDRα4 | DBLδ1 | CIDRβ6 | | | | | |
| PC0053-C.g46 | DBLα0.18 | DBLβ5 | DBLγ10 | DBLδ6 | CIDRβ2 | | | | |
| PC0053-C.g386 | DBLα0.18 | CIDRα4 | DBLβ5 | DBLδ1 | CIDRβ1 | | | | |
| PC0053-C.g671 | DBLα0.18 | CIDRα5 | DBLβ5 | DBLγ10 | | | | | |
| PC0053-C.g689 | DBLα0.18 | CIDRα5 | DBLβ5 | DBLγ13 | | | | | |
| PC0053-C.g585 | DBLα0.2 | CIDRα3.1 | DBLδ1 | CIDRβ5 | | | | | |
| PC0053-C.g612 | DBLα0.2 | CIDRα3.1 | DBLδ1 | CIDRβ1 | | | | | |
| PC0053-C.g682 | DBLα0.21 | CIDRα2.1 | DBLδ1 | CIDRβ5 | | | | | |
| PC0053-C.g656 | DBLα0.22 | CIDRα3.4 | DBLδ1 | CIDRγ11 | | | | | |
| PC0053-C.g268 | DBLα0.3 | CIDRα5 | DBLβ4 | DBLγ18 | DBLε4 | DBLε11 | | | |
| PC0053-C.g410 | DBLα0.3 | CIDRα5 | DBLβ4 | DBLγ13 | DBLζ2 | DBLε4 | | | |
| PC0053-C.g448 | DBLα0.4 | CIDRα6 | DBLβ5 | DBLγ3 | DBLζ4 | | | | |
| PC0053-C.g644 | DBLα0.4 | CIDRα4 | DBLδ1 | CIDRβ1 | | | | | |
| PC0053-C.g421 | DBLα0.5 | CIDRα2.5 | DBLβ13 | DBLδ1 | CIDRβ1 | | | | |
| PC0053-C.g659 | DBLα0.5 | CIDRα2.1 | DBLδ1 | CIDRβ3 | | | | | |
| PC0053-C.g494 | DBLα0.6 | CIDRα3.1 | DBLγ13 | DBLδ1 | CIDRβ1 | | | | |
| PC0053-C.g667 | DBLα0.8 | CIDRα4 | DBLδ1 | CIDRγ9 | | | | | |
| PC0053-C.g474 | DBLα0.9 | CIDRα2.2 | DBLδ1 | CIDRβ1 | | | | | |
| PC0053-C.g535 | DBLα0.9 | CIDRα2.1 | DBLδ1 | CIDRγ4 | | | | | |
| PC0053-C.g556 | DBLα0.9 | CIDRα2.1 | DBLγ10 | DBLδ1 | CIDRγ4 | | | | |
| PC0053-C.g639 | DBLα0.9 | CIDRα2.2 | DBLδ1 | CIDRβ1 | | | | | |
| PC0053-C.g146 | DBLpam1 | DBLpam2 | CIDRpam | DBLpam3 | DBLεpam4 | DBLεpam5 | DBLε10 | | |
| PC0053-C.g30 | DBLδ1 | CIDRγ5 | | | | | | | |
| PC0053-C.g650 | DBLε8 | CIDRα3.2 | DBLδ1 | | | | | | |

New rosetting variant (DC15) - 9197varR1

**Rosetting-associated head structure**

DBLα1.5 /6 /8 | CIDRβ /γ /δ

# PC0053 (9197)
