## Supplementary Figures 2-4 and Supplementary Table1 for "Identification of novel PfEMP1 variants containing domain cassettes 11, 15 and 8 that mediate the *Plasmodium falciparum* virulence-associated rosetting phenotype"

**McLean et al, Supplementary Figures**

**Figure S1. PfEMP1 repertoires of the Kenyan parasite lines.** PfEMP1 domains are colour-coded as described by Rask et al (27). (See separate PDF)

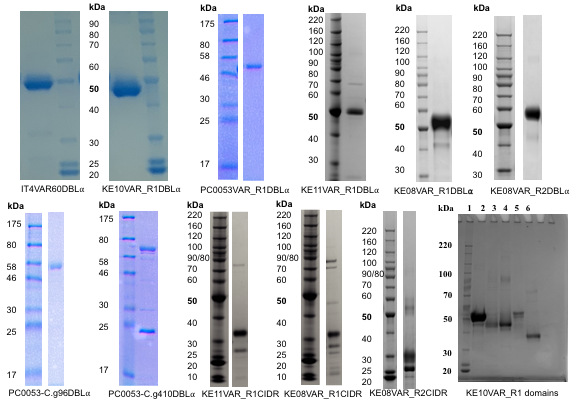

**Figure S2. SDS-PAGE of recombinant PfEMP1 domains.** SDS-PAGE images of the recombinant PfEMP1 domains used to generate antibodies and/or used in erythrocyte binding assays and ELISAs. The his-tagged proteins were expressed in *E. coli* and purified by Ni-NTA or Co-NTA affinity chromatography, followed by size exclusion chromatography in some cases (this was done for all of the DBLα proteins except KE11VAR_R1 and PC0053-C.g410). Proteins IT4VAR60DBLα, KE10VAR_R1DBLα, PC0053VAR_R1DBLα and PC0053-C.g96 had the his-tag removed by TEV cleavage as described previously (17, 48), whereas the other proteins did not. Gels were 4-12% Bis-Tris Novex gels run with MOPS or MES buffer at 200V for 55 minutes and stained with Instant Blue. Domain names are given below each image, except for KE10VAR_R1 domains which are lanes 1) Benchmark ladder (2) PFKE10VAR1 DBLα1.8 (3) PFKE10VAR1 DBLγ7 (4) PFKE10VAR1 DBLε11 (5) PFKE10VAR1 DBLζ3 and (6) PFKE10VAR1 DBLε8. Molecular weight markers in kDa are shown for each gel. The NTS-DBLα preparations gave >95% of protein at the expected molecular weight (~50 kDa), whereas the other domain types had some degraded fragments and/or dimers.

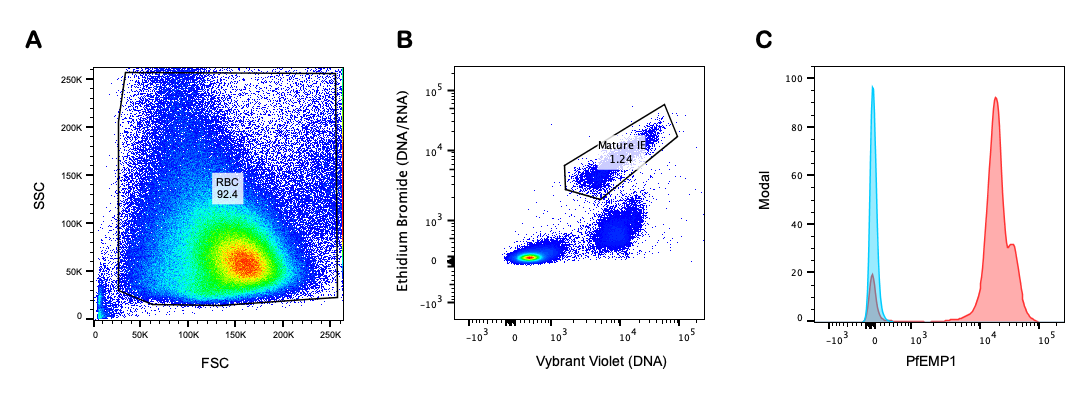

**Figure S3. Gating strategy to identify PfEMP1 on mature infected erythrocytes**. A) Forward and side scatter were used to gate on erythrocytes (RBC) and exclude debris. B) Mature pigmented-trophozoite and schizont infected erythrocytes (IE) were detected as the DNA/RNA high population by staining with 1/2500 dilution of Vybrant DyeCycle Violet (DNA stain) and 20μg/ml of ethidium bromide (DNA/RNA stain). C) PfEMP1 on the surface of live mature infected erythrocytes was detected with 20μg/ml of polyclonal rabbit IgG against NTS-DBLα (variant KE08VAR_R1 shown) followed by 1/1000 dilution of Alexa Fluor 647-conjugated goat anti-rabbit IgG secondary antibody (red). The negative control (blue) was the same parasite culture suspension stained with rabbit IgG against NTS-DBLα from an irrelevant PfEMP1 variant (HB3VAR03 shown) or non-immunised rabbit IgG.

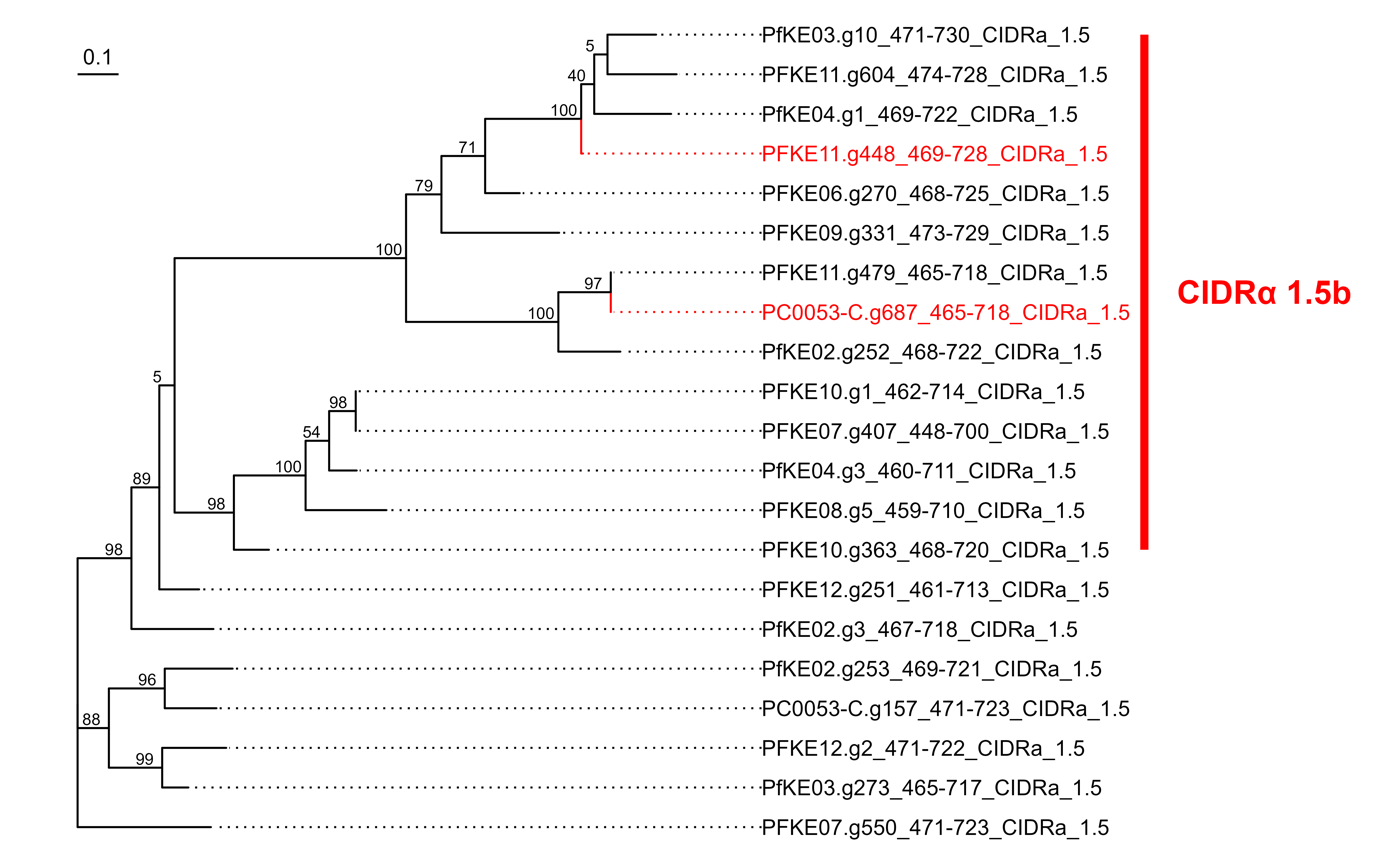

**Figure S4. Phylogenetic tree of CIDRα1.5 domains from the Kenyan parasite lines**. IQTREE was used to generate a maximum likelihood tree of the CIDRα1.5 amino acid sequences from the PFKE parasite line PfEMP1 repertoires, which had been aligned using MUSCLE. The amino acid domain boundaries were obtained from a previous study (34). Percentage bootstrap support is indicated on the nodes based on 1000 replicates, and the scale bar represents the number of changes per site. The CIDRα1.5b domains from the two rosetting variants are indicated in red (PFKE11.g448= PFKE11VAR_R1 and PC0053-C.g687 = PC0053VAR_R1) and are found within a distinct clade within the tree supported by high bootstrap values. In this example, the tree is used as a way of visualising similarity between variants and is not intended to infer evolutionary descent.

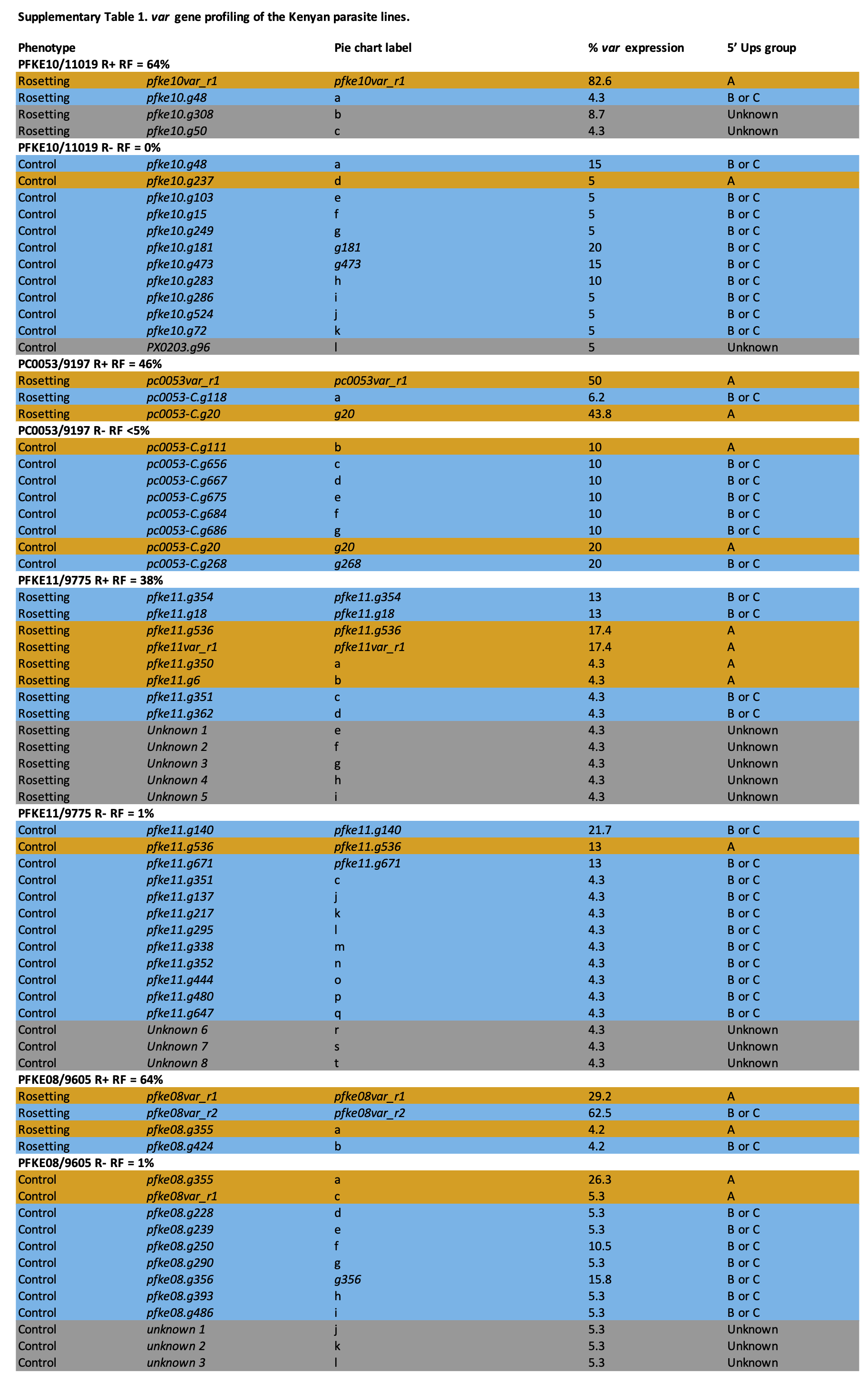
